# Urban soils along the Kern River and Los Gatos Creek are hotspots for *Coccidioides* in the San Joaquin Valley of California

**DOI:** 10.64898/2026.01.15.698504

**Authors:** Robert Wagner, Liliam Montoya, Molly Radosevich, Justin Remais, John W. Taylor

## Abstract

Coccidioidomycosis (Valley fever) is acquired through inhalation of spores produced by fungi in the genus *Coccidioides*. *Coccidioides* is commonly detected in, and cultivated from, the lung tissue of native rodents and the soils within their burrows. Coccidioidomycosis is acquired by exposure to environmentally produced spores and is not spread between hosts. Thus, determining the location of rodent burrows with soils harboring *Coccidioides* will be critical for understanding coccidioidomycosis incidence and modelling how *Coccidioides* distributions will be affected by global change. *Coccidioides* is readily detected in rodent burrows on undisturbed land, has not been detected on agricultural land, and is unstudied on urban land. To test the hypothesis that *Coccidioides* is in urban soil, we sampled rodent burrow soils from the banks of two water ways, the Kern River and Los Gatos Creek, which transect two cities, Bakersfield and Coalinga, in the San Joaquin Valley of California. To test the hypothesis that *Coccidioides* would not be detected at higher elevations, we extended our sampling of rodent burrows along waterways into the mountains of California. From 1178 soil and settled dust samples, we find that *Coccidioides* is found in urban riparian environments in Bakersfield and Coalinga, on riparian land on the floor of the San Joaquin Valley but not at higher elevations and is negatively correlated with modeled soil moisture. *Coccidioides* shows significant co-occurrence patterns with animal-associated fungal taxa, but no broader relationships with the greater fungal community. Our results warrant caution when excavating urban rodent burrows in the region.

**Author Summary:** Coccidioidomycosis, or Valley fever, is caused by a fungus that primarily infects people when its spores become airborne and are inhaled. Although the fungus is found in the burrows of rodents, its presence in cities has been unclear. We investigated whether urban environments in California’s San Joaquin Valley, an area with high Valley fever incidence, harbor the fungus. By sampling more than a thousand soils and settled dust samples along riparian areas (waterways and adjacent environments) in Bakersfield and Coalinga, we found that the fungus that causes Valley fever, *Coccidioides,* occurs in urban riparian zones. The fungus was restricted to low elevations and was more likely to be detected in drier soils. We also compared positive and negative soils using DNA-based surveys of fungi. While the broader fungal community showed no strong relationship to *Coccidioides*, several species tied to animal activity were more common in positive samples, supporting a link to rodent hosts. These findings show that urban waterways can serve as habitat for *Coccidioides*. Because these areas lie within major population centers, identifying and avoiding disturbance of rodent burrows may reduce exposure risk, and future models of Valley fever should account for riparian corridors and local ecological conditions.

## Introduction

*Coccidioides immitis* and *Coccidioides posadasii* are Ascomycete fungi, within the Onygenales, that are responsible for the human disease coccidioidomycosis, colloquially known as “Valley fever” and “Desert Rheumatism” (Posadas 1892, Ophüls 1905, Kollath et al. 2019). Coccidioidomycosis is an important disease in arid regions of the western United States, causing nearly 200 deaths and generating $3.9 billon in healthcare costs per year (Huang et al. 2012, Centers for Disease Control and Prevention 2018, Gorris et al. 2021). In California, Washington and areas generally west of the Sierra Nevada-Cascade mountain ranges, only *C. immitis* has been found, while *C. posadasii* has been observed elsewhere in the United States (Koufopanou et al. 1997, Johnson et al. 2014, Marsden-Haug et al. 2014, Teixeira and Barker 2016, Chow et al. 2021), as well as in Mexico (Baptista-Rosas et al. 2012), Central America (Engelthaler et al. 2016) and South America (Fisher et al. 2001, De Macêdo et al. 2011, Teixeira et al. 2019). Although *Coccidioides* has been found most abundantly in undeveloped arid lands, it remains uncertain whether exposure within cities results from local sources or from travel to nearby rural areas where *Coccidioides* is established. Understanding possible urban exposure is therefore critical for public health, especially in cities such as Bakersfield (population ∼400,000) and Coalinga (∼17,000), which are major population centers in the SJV.

Humans acquire coccidioidomycosis by inhaling arthroconidia from environmental sources (Nguyen et al. 2013) and human-to-human transmission is limited to organ transplantation (Nokes and Blair 2020), which makes determining the geographic location and seasonal timing of arthroconidial development and dispersal important for understanding coccidioidomycosis incidence. All fungi are heterotrophs (organisms that obtain energy and carbon from organic matter rather than producing it themselves), and most fungi, Ascomycota included, consume plants for nutrition. An association between *Coccidioides* spp. and rodents was discovered in the 1940s, suggesting that rodents may serve as a natural reservoir. (Emmons 1942, Emmons et al. 1942). This relationship has been reaffirmed by recent molecular work (Salazar-Hamm 2021, Salazar-Hamm et al. 2022). An explanation for this association comes from genomic analyses showing that *Coccidioides* spp. and closely related genera in the Onygenales evolved away from an ancestral state of using plant carbohydrates for nutrition and to a derived state of using animal protein (Sharpton et al. 2009, Whiston and Taylor 2016). *Coccidioides* spp. are significantly more likely to be found in soil collected from within rodent burrows rather than soil collected apart from rodent burrows (Egeberg and Ely 1956, Elconin et al. 1957, Kollath et al. 2020, Wagner et al. 2023, Head et al. 2024).

There is evidence that *Coccidioides* spp. have evolved a commensal relationship with native rodents, and can persist in the lungs in a dormant state, arrested by the host’s immune system (Sun and Huppert 1976, Rappleye and Goldman 2006, Hsu et al. 2022). This persistence supports the hypothesis that *Coccidioides* functions as an endozoan (occupies an animal host) before reverting to a saprotrophic (decomposer) phase following host death (Taylor and Barker 2019). Fungal endozoans may live as benign parasites in animal hosts without observable symptoms and, following host death, use the host carcass to grow as hyphae and reproduce by arthroconidia.

We previously found that 37% of 238 soils sampled from burrows on undeveloped land in the southwestern San Joaquin Valley (SJV) were positive for *Coccidioides* spp., whereas none of 472 agricultural surface soils sampled from four farms in the SJV were positive for *Coccidioides* spp. (Wagner et al. 2023). There was an observed absence of rodent burrows in agricultural fields at all agricultural sites, likely owing to cultivation, irrigation, and efforts by farmers to eradicate rodents (Baldwin et al. 2016, Lloyd and Baldwin 2021). Unlike undisturbed and agricultural soil, urban soil remains nearly unstudied for *Coccidioides* (Lauer et al. 2012). Having observed that native rodents excavate burrows in the banks of waterways, we hypothesized (H_1_) that burrow soil from urban riparian habitats would harbor *Coccidioides*. Riparian habitats are vegetated corridors bordering rivers or streams that differ sharply from surrounding arid landscapes. Here, we report on the presence of *Coccidioides* in soils sampled from rodent burrows along riparian corridors traversing two cities in the SJV, Bakersfield and Coalinga. Riparian corridors in arid regions have higher plant biomass than surrounding areas (Scott et al. 2014), which provides food for native animals (Case and Kauffman 1997, Pasternak et al. 2013) and increases their abundance and diversity (Patten 1998, Merritt and Bateman 2012, Free et al. 2015). This diversity includes all mammals (Falck et al. 2003, Hamilton et al. 2015), including rodents native to the SJV (Grinnell 1923, Whitford and Kay 1999). Previous research has shown that rodent burrow soils in arid environments host a rich diversity of fungal species (Hawkins 1996, 1999, Wagner et al. 2022), many of which could potentially interact with *Coccidioides*.

Studies of *Coccidioides* in California soils have traditionally focused on the SJV rather than in the adjacent mountains, probably because only one coccidioidomycosis outbreak has been reported at higher elevation (975m), at a Native American midden near Inyokern, California (Plunkett and Swatek 1957). In this previous study, excavated soils within the midden were positive for *Coccidioides*, while soils from outside the midden were negative for *Coccidioides*, as were the lungs of hundreds of native rodents from the area (Plunkett and Swatek 1957). The Kern River and Los Gatos Creek both originate in nearby mountains, which provided us the opportunity to extend our sampling of rodent burrow soils along an elevational gradient. In doing so, we address a knowledge gap about the distribution of *Coccidioides* and challenge a second hypothesis, (H_2_), that *Coccidioides* would not be found in soils sampled from rodent burrows in foothill regions between the SJV floor and elevations as high as 900m.

Most previous research investigating coccidioidomycosis incidence, in the context of landscape environmental variables, has focused on associations with dry, arid soils (Egeberg and Ely 1956, Elconin et al. 1957, Greene et al. 2000, Kollath et al. 2020), and overlooked riparian and riparian-adjacent habitats. Our choice to sample soils from riparian rodent burrows situated in the banks of rivers and streams allowed us to investigate soils that remained relatively dry while being close to water and food resources that support rodent populations. More generally, riparian corridors within the SJV represent some of the last continuous, undeveloped habitats in the region. These waterways bisect large urban areas, potentially situating *Coccidioides* populations in proximity to dense human populations. Finally, only two other studies have compared soil moisture with direct detections of *Coccidioides* in soils (Lauer et al. 2012, Head et al. 2024), allowing us to continue to explore this important knowledge gap.

Here, we survey soil from rodent burrows in two riparian corridors in the SJV, one along the Kern River, which transects Bakersfield, California, and another along Los Gatos Creek, which transects Coalinga, California. We report the presence of *Coccidioides* in terms of location along waterways and, at some locations, seasonality. To understand variation in the presence of *Coccidioides* and the composition of the fungal community, we consider the location of burrows, associations between *Coccidioides* and the soil fungal community, and potential influences of elevation and estimated soil moisture at the time of sampling. Our results help clarify key knowledge gaps about the distribution of *Coccidioides* that may impact the location of arthroconidial development and dispersal. We hope that our research will stimulate research regarding transmission of *Coccidioides* among native rodents and the dispersal of *Coccidioides* spores in ambient air and dust, as well as the inclusion of these aspects of the *Coccidioides* environmental lifecycle into future predictive models regarding spatial and temporal *Coccidioides* distributions.

## Results

1178 samples (soil and dust) were collected and analyzed for the presence of *Coccidioides* using the CocciEnv qPCR assay. Soil samples were collected from 625 rodent burrows from 35 sites along the Kern River, 205 rodent burrows from seven sites in the greater Bakersfield, California area, 248 rodent burrows or adjacent soils from 16 sites along Los Gatos Creek near Coalinga, California, and 100 urban settled dust samples collected from public spaces in Coalinga, California. The soil fungal community was characterized for 68 soil samples (28 from the Kern River and 40 from Los Gatos Creek), by amplicon sequencing of the ITS2 region of fungal ribosomal DNA.

### *Coccidioides* detection

*Coccidioides* was detected in 6.3% (52/830) of rodent burrow soil samples from sites along the Kern River and in and around the urban Bakersfield area and, of these, was detected in 8.3%, 52/625) of rodent burrow soil samples from sites directly along the Kern River and 0% (0/205) of sites in Bakersfield away from the Kern River (S2 Table). From sites along Los Gatos Creek, *Coccidioides* was detected in 23.2% (53/228) of rodent burrow soil samples, 50.0% (10/20) of surface soil samples and in 2% (2/100) of settled dust samples collected from public surfaces in Coalinga (S1 Fig, S2 Table). All surface soil samples were collected from the “Highway 33 West” site at Los Gatos Creek in August 2020, from directly outside the entrances (<1m) from rodent burrows. It was not determined if these surface soils were otherwise associated with the burrows they were collected near. All but two *Coccidioides* positive soil samples from Kern River sites were collected centrally in Bakersfield (Fig 1a), at sites with soil moisture between 0.045 – 0.100 cm^3^/cm^3^ (Fig 1b) and elevations between 98.56 and 192.59 meters (Fig 1c). The other two positive samples were collected from Kern River Parkway West End, not far from the intersection of the Kern River and the California Aqueduct, with soil moisture between 0.058 –0.115 cm^3^/cm^3^ and elevations between 97.26 and 99.86 meters. Of *Coccidioides* positive soil samples from Los Gatos Creek sites, 69.8% (44/63) were collected from areas within 200m of urban or residential land in the town of Coalinga (Fig 2a). The remaining positive samples came from either undeveloped land approximately 10km east of Coalinga (7 positive samples), where soil moisture ranged from 0.052 to 0.136 cm³/cm³ and elevations ranged from 161.72 to 208.48 m (Fig 2a, 2b) or from undeveloped land west of Coalinga (14 positive samples) at elevations below 373 m (Fig 2c, S2 Fig).

**Fig 1.**
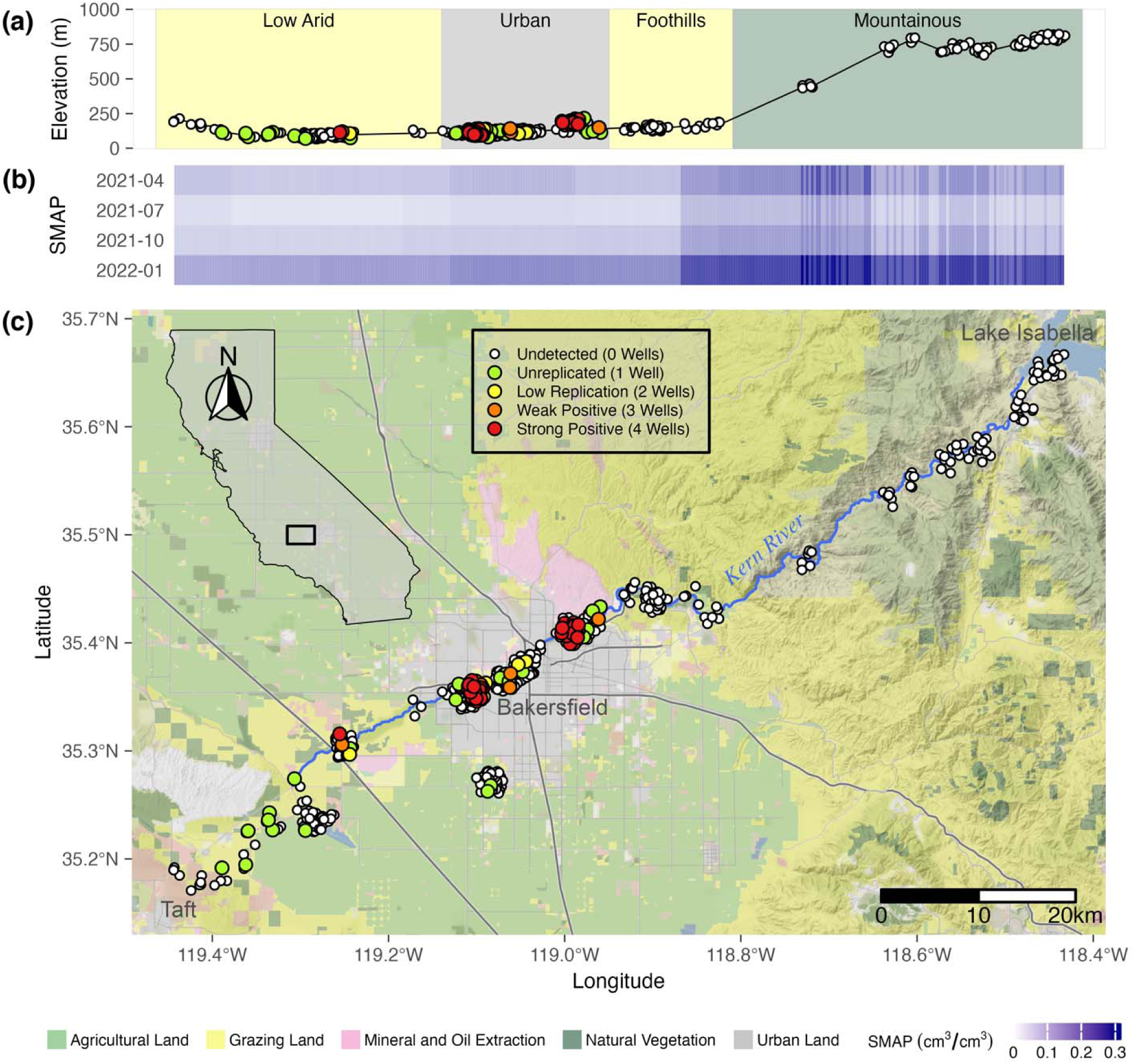
*Coccidioides* detection in rodent burrow soil samples along the Kern River near Bakersfield, California using the CocciENV qPCR assay. Elevation profile (meters above sea level) and qualitative categorization of sample locations (a). Estimated surface soil moisture (upper 5cm) at each timepoint across sampling locations (b). Map of sample coordinates (c). Inset shows map extent. Landscape classification derived from the California Farmland Mapping and Monitoring Program 2018. SMAP = Soil Moisture Active Passive (NASA satellite). Vertical banding in soil moisture is due to varying resolution from sub-9km landscape and vegetation differences, and downscaling algorithms, affecting the SMAP L3 version 3 model. Point colors correspond to positive replicate wells. Map data (lines) were acquired from OpenStreetMap (www.openstreetmap.org) under the Open Data Commons Open Database License (OdbL) (https://www.openstreetmap.org/copyright). Map tiles are by Stamen Design (www.stamen.com) under Creative Commons By Attribution (CC BY 4.0) (https://creativecommons.org/licenses/by/4.0/). California shapefile is public domain data by Natural Earth (http://www.naturalearthdata.com).

**Fig 2.**
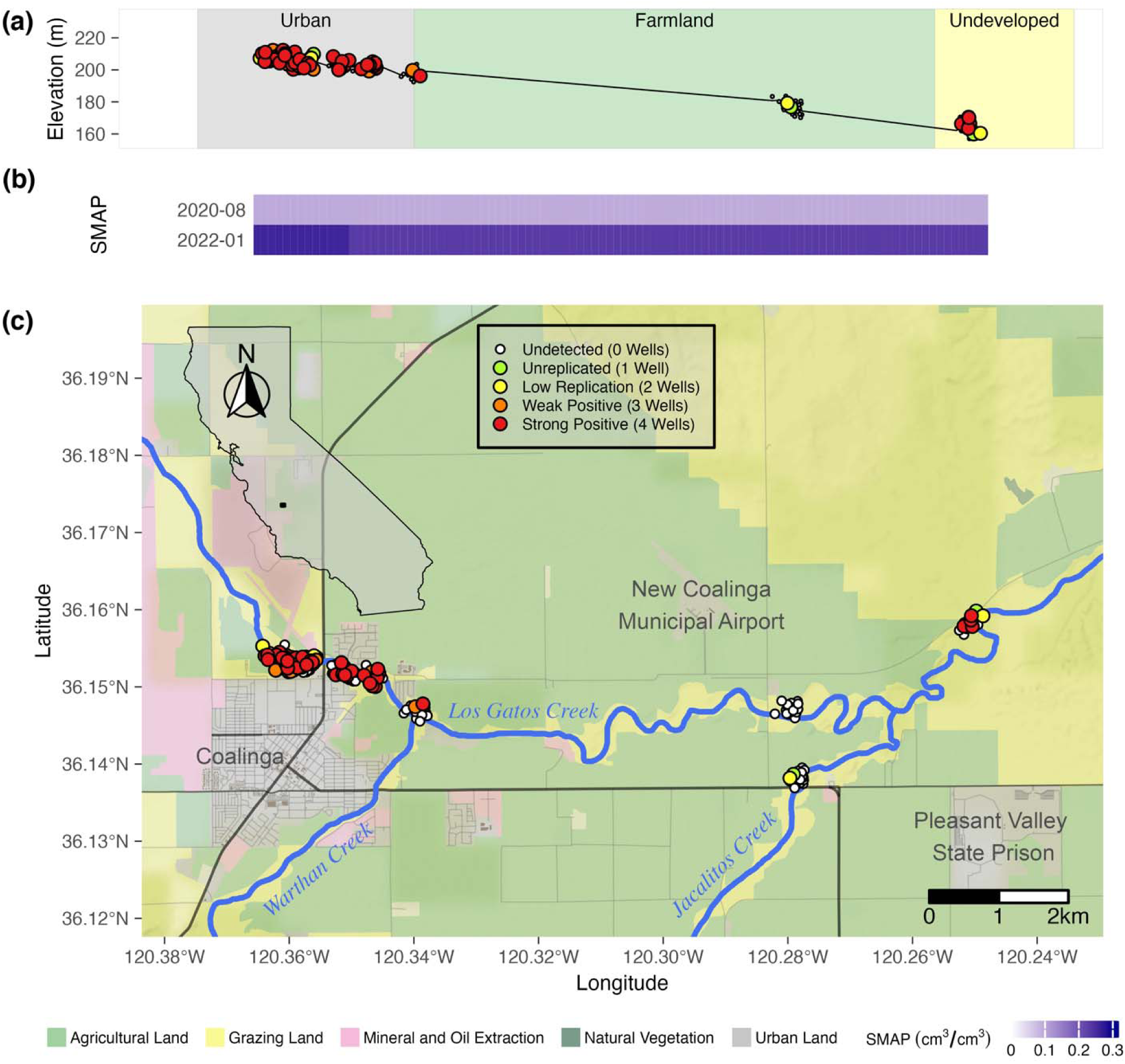
*Coccidioides* detection in rodent burrow and surface soil samples along Los Gatos Creek near Coalinga, California using the CocciENV qPCR assay. Elevation profile (meters above sea level) and qualitative categorization of sample locations (a). Estimated surface soil moisture (upper 5cm) at each timepoint across sampling locations (b). Map of sample coordinates (c). Inset shows map extent. Landscape classification derived from the California Farmland Mapping and Monitoring Program 2018. SMAP = Soil Moisture Active Passive (NASA satellite). Vertical banding in soil moisture is due to sub-9km landscape differences affecting model. Point colors correspond to positive replicate wells. Map data (lines) were acquired from OpenStreetMap (www.openstreetmap.org) under the Open Data Commons Open Database License (OdbL) (https://www.openstreetmap.org/copyright). Map tiles are by Stamen Design (www.stamen.com) under Creative Commons By Attribution (CC BY 4.0) (https://creativecommons.org/licenses/by/4.0/). California shapefile is public domain data by Natural Earth (www.naturalearthdata.com).

Of the 35 sites investigated along the Kern River (ignoring non-Kern River sites), more than 80% of samples positive for *Coccidioides* (45/52) were collected from two sites: University Place (25/106 positive) and Panorama Park West (20/96 positive) (Fig 3a, b). Of the 16 sites investigated along Los Gatos Creek, three sites provided the bulk of *Coccidioides* positive samples: Highway 33 West (27/80 positive), Highway 33 East (15/40 positive), and Phelps Avenue-10km (5/20 positive) (Fig 3c, d). Analyses including soil moisture were limited to samples from Kern River sites. Across all Kern River sites, *Coccidioides* detection showed a moderately significant association with sampling timepoint (p = 0.046, 0.039, 0.040, OR = 0.02, 0.02, 2.52E+03) and surface soil moisture (p = 0.049, OR = 0.06), and no significant association with rootzone soil moisture (p = 0.104) or elevation (p = 0.076) (S5 Table). When excluding samples collected above 210 meters, which excludes foothill sites east and upland of the Kern River Canyon entrance, *Coccidioides* detection did not show significant associations with sampling timepoint, surface moisture and rootzone moisture. This result indicates possible collinearity between these terms, with higher elevations (within the context of lower elevation sites) being a significant predictor (p = 0.014, OR = 3.70E+03) of *Coccidioides* detection (S6 Table). Taken together, *Coccidioides* presence did not show any strong or unambiguous seasonality. Surface soil moisture along the Kern River, increased from west to east, from low to high elevations, and during spring (April 2021) and winter (January 2022), as compared to summer (2021) and fall (2021) (Fig 1b). Along Los Gatos Creek, soil moisture was higher in winter than during the fall (Fig 2b). Despite including site as a random effect to account for site-to-site variability, it’s important to note that 87.5% of all Kern River positive detections for *Coccidioides* occurred at two sites, University Place and Panorama Park West, and that comparing only these two sites showed no significant associations between *Coccidioides* detection and site, sampling timepoint or soil moisture (Fig 4, S7 Table).

**Fig 3.**
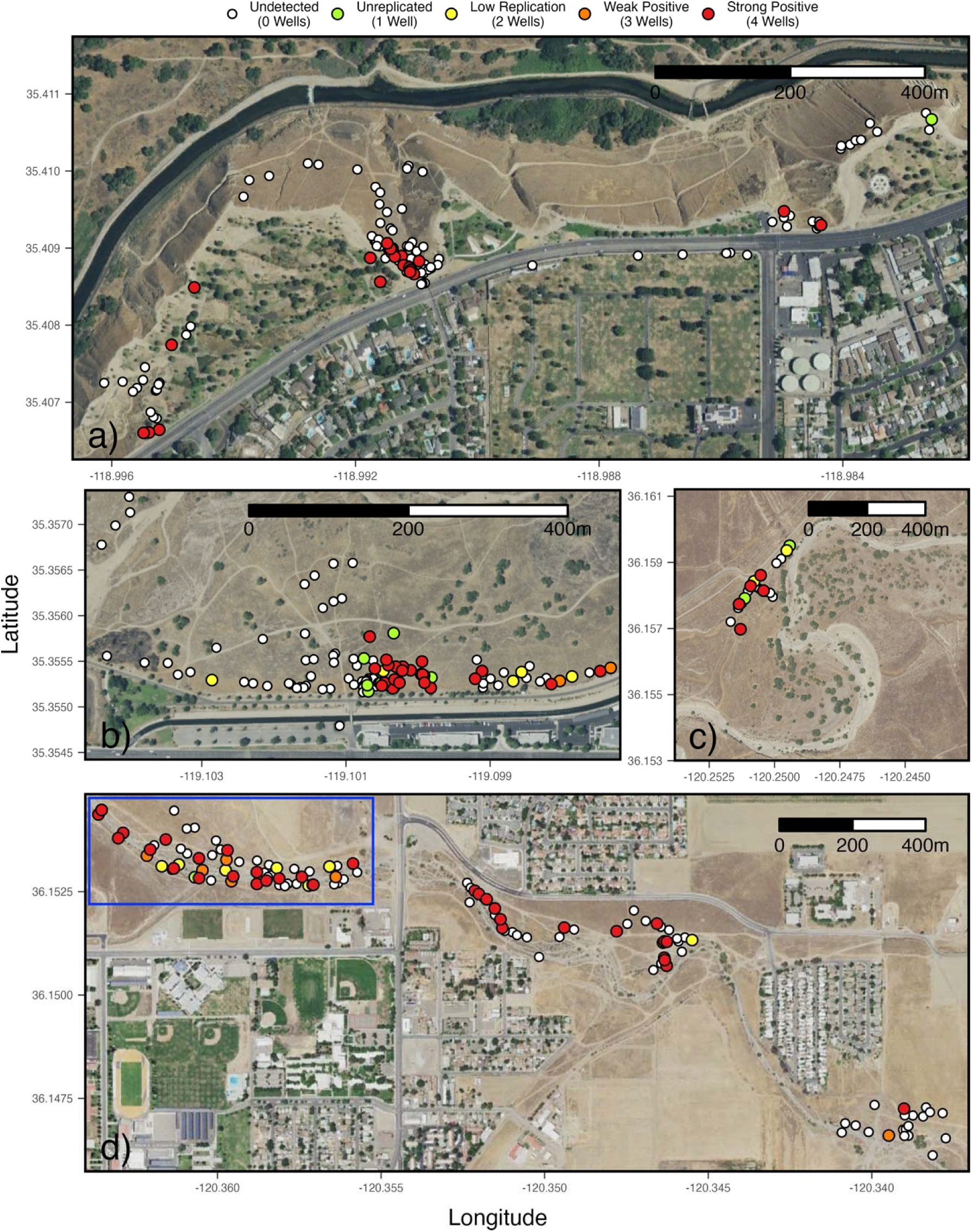
*Coccidioides* detection hotspots. Panorama Park West and Panorama Park East, Kern River (a). University Place, Kern River (b). Phelps Avenue 10km, Lost Gatos Creek (c). Highway 33 West (in blue rectangle, includes 20 surface samples at the same points as burrow samples), Highway 33 East and Warthan Creek, Los Gatos Creek (d). All points jittered to avoid overlapping. Satellite imagery is public domain data provided by the United States Department of Agriculture (USDA) National Agriculture Imagery Program (NAIP) (https://naip-usdaonline.hub.arcgis.com).

**Fig 4.**
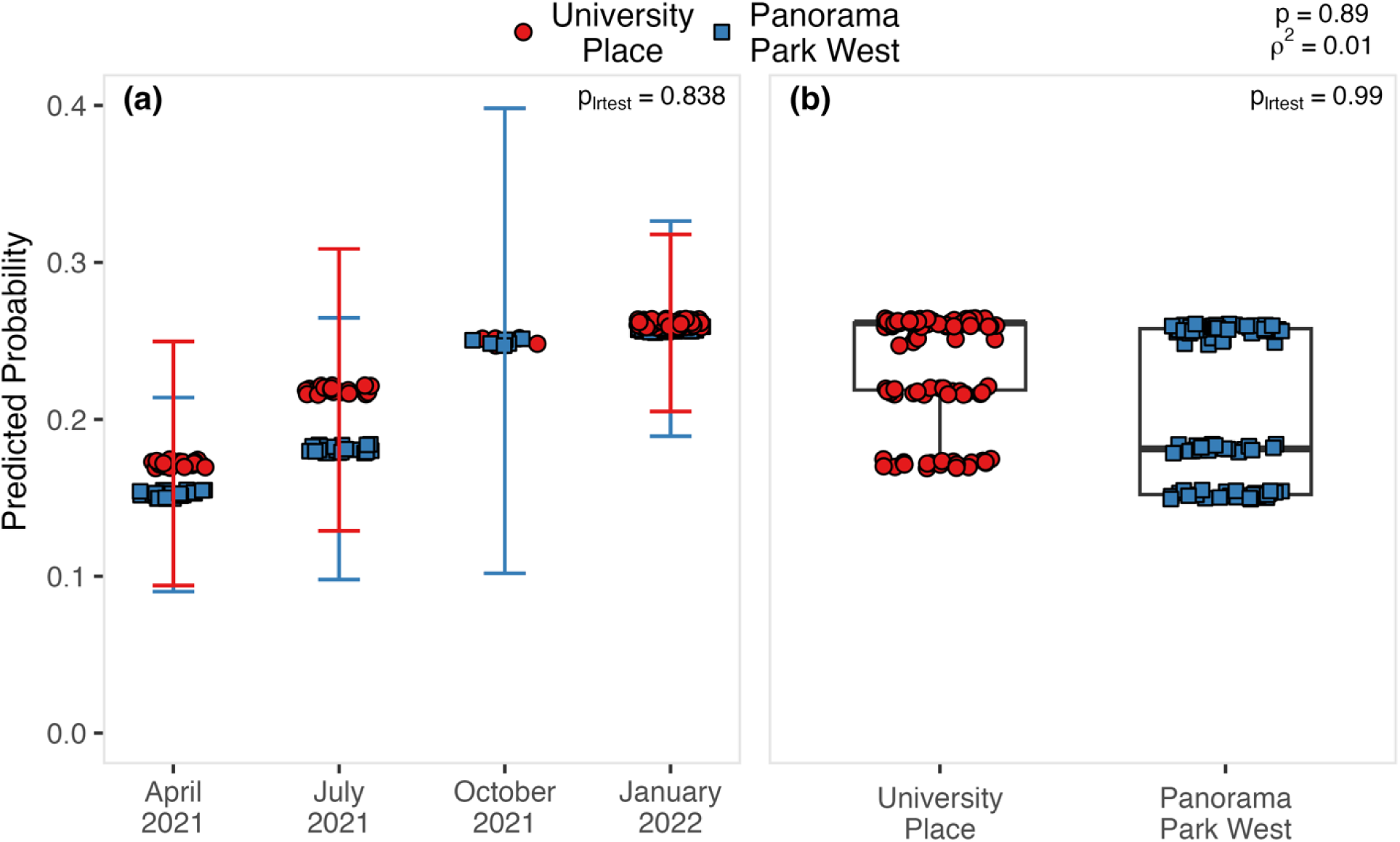
The predicted probability of detecting *Coccidioides* in rodent burrows as a function of timepoint and site (University Place vs Panorama Park West). Mean predicted probabilities are displayed as a function of timepoint (a) and as a function of site (b). Points jittered to show sampling depth. P_lrtest_ = significance of difference with null model lacking a timepoint variable (a) or a site variable (b) derived from a log-ratio test. “rho squared” (ρ^2^) = McFadden’s-pseudo-R^2^ for full logistic regression model. p = significance of full logistic regression model derived from deviance and null deviance. n = 202. Site error bar for October 2021 perfectly overlap.

### *Coccidioides* and the soil fungal community

For Kern River samples, a significant (p = 0.002) association was found between the structure of the soil fungal community and the site where soils were collected, representing 22% (r^2^ = 0.22) of the variance in Bray-Curtis dissimilarity among all identified fungal species (S8 Table). A post-hoc, pairwise comparison showed a significant difference (p = 0.04, r^2^ = 0.08) between fungal communities from rodent burrow soil samples taken from Panorama Park West and University Place (S9 Table), which, together, account for 23 of the 28 Kern River samples sequenced for ITS2. However, when examining all sites, no significant associations were found between the structure of the soil fungal community (Bray-Curtis dissimilarity) and the presence of *Coccidioides* in soils. This result was true for rodent burrow samples collected from along the Kern River (Fig 5a, S8 Table), as well as soils collected from within or just outside the entrance of rodent burrows from Los Gatos Creek (Fig 5b, S10 Table). Significant correlations between variables of interest and Bray-Curtis dissimilarity, and corresponding clustering on ordination plots (of which we did not find here), are indicative of ecological relationships (Bray and Curtis 1957, Legendre and Legendre 2012). We found no such correlations here, which does not preclude, but fails to provide evidence supporting an ecological relationship between the fungal community and *Coccidioides*. It is notable that, within the examined fungal communities, species of one genus, *Alternaria*, were most abundant among samples from both Kern River (14.6 ± 2.0%) and Los Gatos Creek (15.4 ± 1.7%) (S3 Fig, S4 Fig). *Alternaria* spp, were also seen to dominate rodent burrow fungal communities in our previous study of sites along California Highway 33 (Wagner et al. 2022). Other abundant genera were, from Kern River samples, *Cladosporium* (11.5 ± 1.6%), *Trichophaeopsis* (6.5 ± 126%), *Neosetophoma* (5.1 ± 1.3%), and *Penicillium* (4.6 ± 0.8%) and, from Los Gatos Creek samples, *Coniochaeta* (7.6 ± 1.8%), *Penicillium* (7.4 ± 1.6%), *Cladosporium* (5.8 ± 0.9%), and *Chaetomium* (2.8 ± 0.6%) (S11 Table).

**Fig 5.**
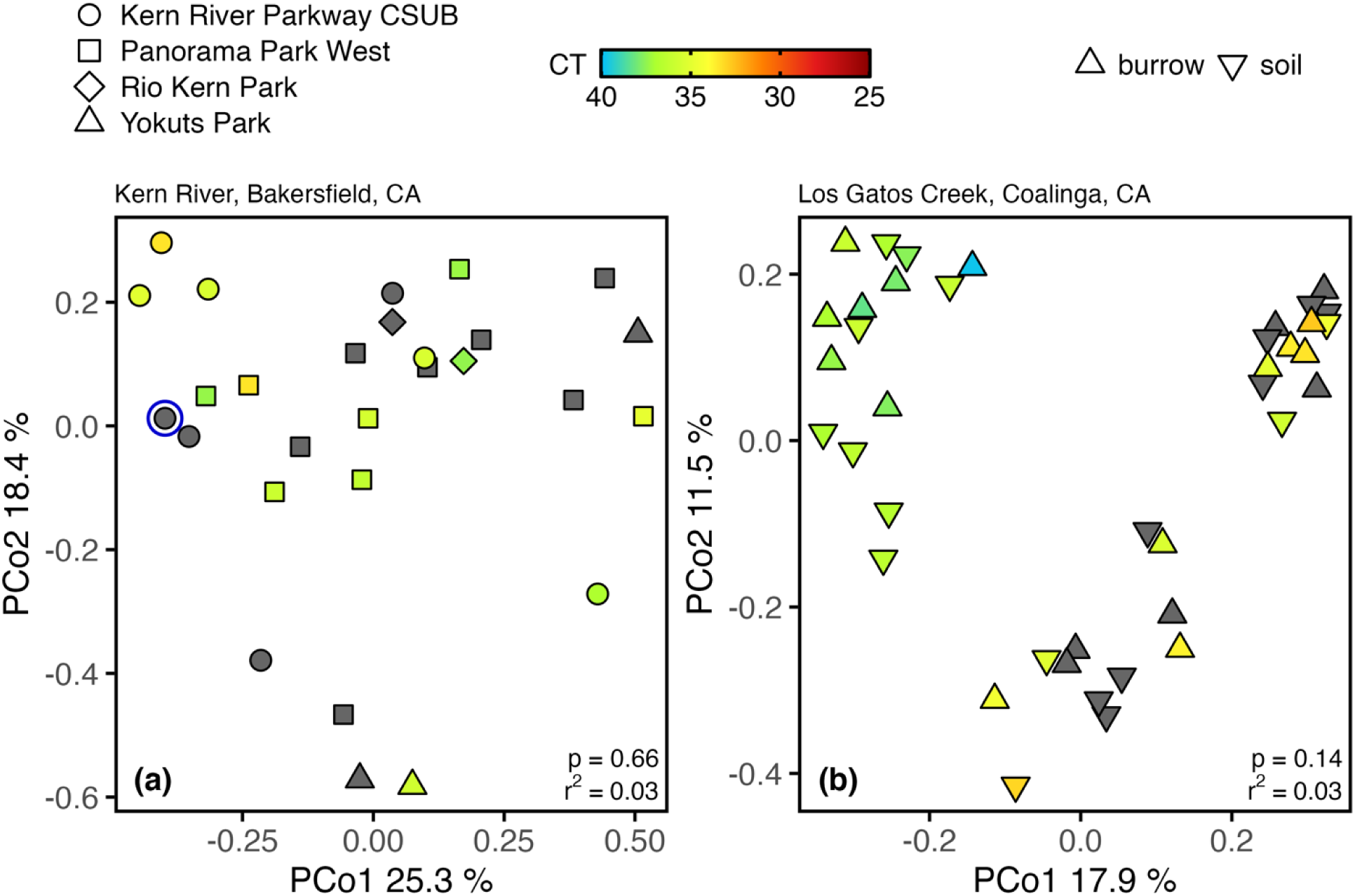
PCoA plot showing the Bray-Curtis dissimilarity within the fungal community between individual rodent burrow soil samples from Kern River sites in Bakersfield, CA (a) and Los Gatos Creek in Coalinga, CA (b). Kern River samples were collected from within rodent burrows in April and July 2021. Los Gatos Creek samples were collected just outside (Entrance) and within (Interior) rodent burrows in August 2020. Colored points = samples positive for *Coccidioides* using the CocciENV qPCR assay. Gray points = samples negative for *Coccidioides* using the CocciENV qPCR assay. Blue circle = sample positive for *Coccidioides* in the ITS2 rDNA dataset. CT = qPCR cycle threshold value. n = 28 (Kern River) and 40 (Los Gatos Creek).

### Indicator species

From the 68 fungal communities sequenced from soils, in which a total of 764 unique fungal species were detected, 83 fungal indicator species (p ≤ 0.05) were identified, 21 with *Coccidioides* positive soils and 62 with *Coccidioides* negative soils. Soils from rodent burrow interiors yielded 11 positive and seven negative associations from Kern River samples (S12 Table) and seven positive and 51 negative associations from Los Gatos Creek samples (S13 Table, 14 Table). Soils collected from outside rodent burrow entrances along Los Gatos Creek provided three positive and four negative associations (S15 Table). The most significant (p ≤ 0.01) associations with *Coccidioides* positive rodent burrow soils were found with *Coniochaeta discospora* and *Montagnea arenaria* from Kern River samples and *Coniochaeta polymorpha* from Los Gatos Creek samples. No similarly significant associations were found with *Coccidioides* negative rodent burrow soils from Kern River samples. Among Los Gatos Creek samples, negative associations with *Aspergillus* spp., *Chaetomium* spp., *Holtermanniella* spp., *Ustilago bullata*, *Paraphoma fimeti*, *Septoriella neoarundinis* and *Pseudoechria decidua* were found for soils within rodent burrows, and with *Dothiora viticola* for soils from outside rodent burrow entrances, at the same (p ≤ 0.01) significance level. A single species, *Fusarium solani*, was identified as an indicator species from both Kern River (p = 0.03) and Los Gatos Creek (p = 0.03) samples, and it was associated with *Coccidioides* negative samples.

## Discussion

Our most important finding is that *Coccidioides* can be found in soils along seasonal waterways in major population centers of the San Joaquin Valley; Bakersfield (population ∼400,000) and Coalinga (∼17,000), where Valley fever incidence is high. This result was true for an ephemeral stream on the west side of the SJV, Los Gatos Creek, in Coalinga, CA and for a seasonally wet, major river, the Kern River, that runs east to west through Bakersfield, CA. We rarely found *Coccidioides* in urban areas away from a waterway, though our sampling in these areas was more limited. In Coalinga, *Coccidioides* was found in only two of the 100 settled dust samples we collected, 0.6 – 3.2 km from Los Gatos Creek. At the Sports Village Complex in Bakersfield, CA, 13 km from the Kern River, none of 87 soils samples taken quarterly over one year was positive for *Coccidioides*. Taken together, these results fail to disprove H_1_, that *Coccidioides* would be present in urban riparian areas. The inverse, that *Coccidioides* is absent from non-riparian urban areas cannot be claimed because sampling intensity and site selection may have influenced detection. The close juxtaposition of soils positive and negative for *Coccidioides* within sites where the fungus was commonly detected, e.g., University Place and Panorama Park West (Fig 3b), suggests that *Coccidioides* distribution may not be uniform, consistent with previous reports of sporadic detection in soils (Maddy 1965, Greene et al. 2000, Baptista-Rosas et al. 2012, Litvintseva et al. 2015). While *Coccidioides* is associated with rodents and their burrows in the SJV (Egeberg and Ely 1956, Elconin et al. 1957, Kollath et al. 2020), our study does not address *Coccidioides* dispersal between rodent burrows or among rodent species.

We found a higher probability of detecting *Coccidioides* in drier soils, though this result is almost certainly tied to other variables such as elevation and site choices. Previous research has suggested that dry soils are associated with coccidioidomycosis incidence, potentially due to competition between *Coccidioides* and other soil microbiota (Cordeiro et al. 2006, Kollath et al. 2019, Dobos et al. 2021, Head et al. 2024, Porter et al. 2024). Among our Kern River samples, no *Coccidioides* positive samples were collected from sites with monthly average soil moisture above 10% v/v. The higher soil moisture levels associated with drainage and concentrated water flow in the Kern River Canyon may contribute to the absence of *Coccidioides* in soils at higher elevations in our study. Downstream areas, where the Kern River is ephemeral due to agricultural water diversion and flows only during high rainfall years, showed greater prevalence, suggesting that relatively lower soil moisture may facilitate *Coccidioides* presence. However, regions with extremely low soil moisture, such as between Bakersfield and Taft, presented limited detection, possibly due to reduced availability of vegetation and water to support rodent hosts. These observations suggest that soil moisture may influence *Coccidioides* presence, though other factors are likely involved. Epidemiologic studies have shown that in typically arid regions, periods of above-average rainfall are often followed by increased coccidioidomycosis incidence several months to a year later, likely due to enhanced fungal growth during moist conditions and subsequent spore release as soils dry (Comrie 2005, Park et al. 2005, Tamerius and Comrie 2011, Head et al. 2022, Kollath et al. 2022). This pattern supports the idea that *Coccidioides* benefits from intermittent or moderate moisture, but not persistently wet conditions. Our conclusions regarding *Coccidioides* population dynamics relative to soil moisture are limited by 1) the coarse, 9 km^2^ grid scale resolution of modelled soil moisture data, 2) the fact that most positive samples came from only two of the 42 Kern River and greater Bakersfield sites and 3) covariation of soil moisture with elevation. Despite these limitations, our results provide a landscape-level comparison of *Coccidioides* presence with modeled soil moisture, contextualizing previous work linking soil moisture and coccidioidomycosis incidence (Gorris et al. 2018).

A third observation is the absence of *Coccidioides* at elevations between 373 and 888 m, with most positive detections below 250 m. This result fails to disprove H_2_, that *Coccidioides* would not be detected in foothill and montane sites and is likely influenced by soil moisture and other environmental variables that correlate with elevation. We commonly observed the diurnal California ground squirrel (*Otospermophilus beecheyi*) upslope at foothill and montane sites, similar to our observations at low elevation sites on the valley floor. Deer mouse (*Peromyscus maniculatus*) and San Joaquin pocket mouse (*Perognathus inornatus*), rodent species known to harbor *Coccidioides* (Emmons 1942, Catalán-Dibene et al. 2014), are reported to inhabit sites from Bakersfield to our highest sampling sites at Lake Isabella (Best 1993, Ocampo-Chavira et al. 2020, Laabs et al. 2022). Thus, observed rodent host populations at higher elevations suggest that host presence or absence alone cannot explain the absence of *Coccidioides* at these sites.

### Fungal Community Characteristics

We did not find consistent associations between *Coccidioides* and overall soil fungal community structure, consistent with both our previous research along California Highway 33, 60 km west of Bakersfield, CA (Wagner et al. 2022) and more recent work done in the Carrizo Plain (Radosevich et al. 2024). Our use of nucleic acids for identification does not distinguish between dormant spores and actively growing mycelia, nor between living and dead material, limiting inferences about community interactions. An abundance of spores relative to mycelia could bias our understanding of the soil fungal community (Nilsson et al. 2019). Additional research will be needed to determine if meaningful interactions between *Coccidioides* and the broader soil fungal community exist. Low soil moisture may correlate with spore abundances in arid soils (Liu et al. 2009, Deveautour et al. 2020), and dry conditions can limit fungal growth (Ramirez et al. 2004, Magan 2007). We did not directly measure soil moisture content, though we qualitatively observed that the soils we sampled were dry.

Indicator species analysis identified several individual fungi associated with *Coccidioides* positive samples, including *Coniochaeta discospora* and *Coniochaeta polymorpha*, as well as *Preussia spp.*, *Gymnoascus dugwayensis*, and *Myxotrichum deflexum*. Many of these taxa are coprophilous or associated with animal-derived substrates, suggesting a potential link between *Coccidioides* presence and rodent activity. This finding agrees with recent research showing that rodent presence and soil *Coccidioides* presence are correlated in and around giant kangaroo rat (*Dipodomys ingens*) burrows in the nearby Carrizo Plain (Head et al. 2024).

*Coniochaeta* spp. are soil saprotrophic and animal pathogenic fungi, of which many species are coprophilous (live on animal dung) (Richardson 1998, Lytvynenko and Hayova 2018). There is limited evidence that *Coniochaeta* species are keratinophilic (consumes keratin), having been isolated from desert soil crusts using sheep wool enriched media (Hamm et al. 2020). *Coniochaeta discospora* is commonly found on mammal dung (Cain 1957, Hanks 1963, Mahoney and LaFavre 1981), while *Coniochaeta polymorpha* is a more recently identified species isolated from human tracheal (Khan et al. 2013) and gastric (Heydari et al. 2021) samples, and that morphologically resembles *Coniochaeta hoffmannii*, an opportunistic pathogen of humans, dogs and cattle (Sakaeyama et al. 2007, Hurst 2019). Some *Coniochaeta* species, including close relatives of *C. discospora*, are also known as opportunistic wood and tree pathogens in Prunus and other hosts (Damm et al. 2010), possibly connecting the fungal communities identified here with the riparian habitats they were isolated from. Most *Preussia* species are coprophilous (Ahmed and Cain 1972), while *Gymnoascus* spp. are within the Onygenales (which include many taxa known to degrade animal protein, including *Coccidioides* spp.), produce keratinase and have been isolated from hair and wool (Błyskal 2009). *Myxotrichum* spp., also within the Onygenales, are soil saprotrophs that are mostly associated with breaking down plant cellulose, though also produce enzymes that can degrade animal tissue (Rice 2005). Our data do not establish a causal relationship, but may indicate an association between *Coccidioides* positive samples and actively inhabited rodent burrows, concurring with research done in the Carrizo Plain of California (Head et al. 2024).

### *Coccidioides* biogeography

Landscape models predict *Coccidioides* distribution based on soil, climate, and case data (Gorris et al. 2019, Weaver et al. 2020, Dobos et al. 2021). However, no current landscape modeling efforts take into consideration environmental detections or rodent host ecology. Our findings of sporadic detection in riparian soils, and our previous finding of absence in agricultural soils (Wagner et al. 2023) refine understanding of habitat suitability and suggest that water availability and local environmental conditions may indirectly influence *Coccidioides* presence. Namely, by suggesting that nearby water resources can influence the distribution of *Coccidioides* in rodent burrow soils through the provision of ecosystem resources, either directly, or by supporting local rodent populations (Falck et al. 2003, Hamilton et al. 2015). Indeed, early work isolated *Coccidioides* from sites in the SJV along both the Fresno and Chowchilla Rivers in Merced and Madera counties (Lacy and Swatek 1974). Taking the geography of riparian corridors into account could influence future modeling efforts regarding the predicted landscape-level distribution of *Coccidioides* in the environment and add important constraints to modeling efforts that rely primarily on coccidioidomycosis incidence data. The inclusion of species distribution modelling, focused on likely rodent reservoirs, can further improve such modelling approaches.

### Temporal dynamics and dispersal

No clear temporal trends were detected in rodent burrow soil samples at University Place and Panorama Park West, which accounted for most positive Kern River samples. Limited sampling intervals, presence-absence data, and uneven spatial distribution constrain inferences about seasonality or growth dynamics. Our finding a significant association between *Coccidioides* presence and sampling timepoint across all Kern River samples may indicate unmeasured categorical differences between sites where *Coccidioides* is heavily established, and sites where *Coccidioides* may be limited or more recently established.

*Coccidioides* appears to be distributed non-uniformly across urban riparian landscapes, consistent with prior studies (Maddy 1965, Lacy and Swatek 1974, Greene et al. 2000). Detection at only a subset of sites with similar environmental features (numerous rodent burrows, observably large populations of *O. beecheyi*, irrigated lawns, proximity to the Kern River and close contact with humans) highlights unmeasured variables likely influencing distribution. Patterns may be influenced by rodent populations or localized microhabitat conditions, though these cannot be confirmed here. The distribution of plague, another microbial, zoonotic disease in California associated with *O. beecheyi* (Eskey and Haas 1940, Lang 2004), is better explained by rodent host populations than by climatic niche data (Fell et al. 2022). The possibility that *Coccidioides* has a non-uniform geographic distribution associated with native rodents, not unlike distribution patterns found in plague, suggests improvements for future landscape-level modelling efforts.

### Additional methodological limitations

Although we obtained 1178 samples, sample size and the limit of detection remain the two biggest limitations of our approach (Wagner et al. 2022, 2023). Thus, our inability to detect *Coccidioides* at certain sites does not necessarily preclude the presence of *Coccidioides* at those sites. This limitation is ever greater for settled dust owing to the far smaller size of settled dust samples compared to soil samples. DNA can come from living or dead fungi and may confound our interpretation of temporal dynamics. We know that there is a lower limit of detection for the CocciENV qPCR assay (Bowers et al. 2019), so *Coccidioides* could be present in samples that we scored as absent (Dineen et al. 2010). Additionally, consideration must be given to unknown environmental influences that may affect our ability to detect *Coccidioides* in such samples. Studies of total bacterial and fungal community dynamics at continental or regional scales (Barberán et al. 2015) have successfully used approaches similar to ours (Wagner et al. 2023), but understanding the relationships between *Coccidioides* and individual microbial species may benefit from a targeted approach that investigates a more limited number of fungal taxa. Due to finite sampling resources, nearly all our samples are from within rodent burrows, as this was the most straight-forward way to determine if *Coccidioides* was in an area. However, this approach is unable to address questions regarding *Coccidioides* in other environments. Because *Coccidioides* occurs patchily in soil, we prioritized broad spatial coverage across ecological gradients, complemented by more intensive resampling at sites where detections were frequent. This design provided both regional context and local replication for temporal analyses. While our effort exceeds the sampling scale of most prior environmental surveys (S1 Table), we recognize that further sampling would help quantify detection variability and additional balanced sampling across regions will be needed to confirm observed trends. Although rodent burrows were a focal sampling microhabitat, we did not formally collect data on rodent abundance, activity, or species composition. This study was framed from a microbial ecology perspective to characterize environmental *Coccidioides* distribution patterns. However, the close ecological association between *Coccidioides* and rodent hosts suggests that integrating host ecology will be essential for defining mechanistic interactions. We observed a notably high number of *O. beecheyi* at several Kern River sites, especially at University Place and Panorama Park West. Dozens were regularly seen during fieldwork, often retreating into the same burrows we sampled. *O. beecheyi* have been associated with *Coccidioides* presence in Mojave Desert soils (Lauer et al. 2020), and related rodents have tested positive for *Coccidioides* in lung tissue (Emmons et al. 1942, Salazar-Hamm 2021). Many sampled burrows showed small mammal footprints consistent with active habitation (Grinnell 1923, McNew et al. 2023). While our observations suggest a potential link between *O. beecheyi* behavior and *Coccidioides* distribution, it is important to emphasize that future testing of *O. beecheyi* tissues (Baptista-Rosas et al. 2012, Salazar-Hamm 2021) is needed to confirm of refute their potential role in *Coccidioides* ecology. Our investigation of riparian zones in urban population centers means that our samples largely came from semi-natural areas situated in urban centers, making land classification challenging. Finally, our use of 9km^2^ grid scale soil moisture data, derived from satellite measurements, limits the conclusions we can draw regarding comparisons between *Coccidioides*, the soil fungal community and soil moisture levels. Future studies should pair direct soil moisture measurements with molecular surveys for *Coccidioides* and other fungal species.

### Conclusion

Our findings, based on an analysis of 1178 samples, revealed that *Coccidioides* is abundant, though unevenly distributed, in soils along two waterways that cross major urban population centers in the SJV, a region high in coccidioidomycosis incidence. The distribution pattern of *Coccidioides* showcases a clear elevation-dependent dynamic, suggesting that elevation, environmental variables that correlate with elevation, or both, play a pivotal role in shaping *Coccidioides* biogeography. Our samples were primarily obtained from rodent burrows, and possible constraints on transmission among rodent groups are suggested by the proximity of *Coccidioides* positive and negative soils at single sites throughout the year. Our investigation into the relationship between *Coccidioides* and the broader soil fungal community yielded consistent results with our previous study along Highway 33, with both providing no evidence for strong, community-level associations, challenging assumptions about the activity of *Coccidioides* outside of animal hosts. However, our identification of a handful of individual fungal taxa that correlated with *Coccidioides* presence, most of which live on animal derived substrates, provides evidence that *Coccidioides* presence may be associated with rodent activity. Finally, the identification of riparian corridors as potential *Coccidioides* habitats emphasizes the need for nuanced geographical considerations in predictive models, taking into consideration both landscape topography, landscape use, and distributions of rodent hosts.

Our detection of *Coccidioides* in native rodent burrows situated within and adjacent to major population centers in the SJV underscores a previously underappreciated risk to public health. The highest incidence of Coccidioidomycosis in California is in Kern County (Nguyen et al. 2019), nearly half the population of which resides in the city of Bakersfield, where we found abundant *Coccidioides*. Likewise, Coalinga is the largest population center in western Fresno County, where we also regularly found *Coccidioides*. Waterways, likely home to potential *Coccidioides* rodent hosts, bisect or run adjacent to many other population centers in the SJV.

These findings advocate for similar surveys in population centers elsewhere in the SJV and for targeted mitigation strategies that prioritize the identification and non-disturbance of rodent burrows and nearby soils. Where disturbance is unavoidable, stringent adherence to personal protective protocols is essential (Das et al. 2012). Importantly, efforts to exterminate native rodents may prove counterproductive; the resultant increase in carcasses could enhance fungal growth and sporulation, thereby exacerbating the incidence of Valley fever in the area. Similarly, the effectiveness of trapping and removing rodents is uncertain given the extensive reservoir of susceptible rodents in cities and surrounding peri-urban landscapes. Preventing access to discarded human food (and access to other human-derived food resources, i.e. irrigated lawns) may provide a more sustainable and long-term option for limiting potential *Coccidioides* rodent host populations. Collectively, our results highlight the need for integrated management strategies that address both the ecological complexity of *Coccidioides* and public health imperatives.

## Materials and Methods

We sampled soil at 59 individual sites in and around the cities of Bakersfield and Coalinga in California’s SJV (Fig 1), with the intent of obtaining latitudinal transects describing *Coccidioides* presence across both population centers. Because they are accessible, create natural uninterrupted transects across landscapes and support native rodent populations whose burrows are associated with *Coccidioides*, sampling occurred primarily along two seasonally ephemeral waterways, the Kern River in Kern County, and Los Gatos Creek in Fresno County. The Kern River originates in the Sierra Nevada mountains in Sequoia National Park (United States Geological Survey 2021a), travels south to a reservoir, Lake Isabella, and then west through Bakersfield before terminating at the California Aqueduct near the Buena Vista Aquatic Recreational Area in the southern SJV. Los Gatos Creek begins in the Clear Creek Management Area in California’s Diablo Range (United States Geological Survey 2021b), flows east through Coalinga and terminates at the California Aqueduct near Huron, approximately 130km north of the Kern River terminus. Soils in the vicinity of our study sites along both waterways are aridic and receive little annual rainfall. Wild vegetation along Los Gatos Creek is dominated by tree species of *Quercus*, *Acer* and *Pinus* at higher elevations and at lower elevations by grass/shrub species of *Nassella, Sporobolus,* and *Suaeda nigra* (Griffith et al. 2016). Wild vegetation along the Kern River is classified in three ecoregions: one characterized by *Allenrolfea occidentalis*, *Atriplex spp*., *Nassella spp*. and *Prosopis spp*. from Taft to Bakersfield; a second characterized by *Atriplex spp*., *Nassella* spp., and some *Quercus douglasii* and *Opuntia basilaris* from Bakersfield to the Sierra foothills; and a third characterized by *Adenostoma fasciculatum*, *Ceanothus spp.*, *Nassella spp.*, *Pinus sabiniana* and *Quercus spp.* found east from the Sierra foothills along an increasing elevation gradient to Lake Isabella (Griffith et al. 2016).

### Sampling and DNA Extraction

We collected soil from 625 rodent burrows at 35 sites along the Kern River and from an additional 205 rodent burrows at seven sites away from waterways (non-Kern River sites) in and around Bakersfield and Taft, California. In Coalinga we collected soils from 228 rodent burrows and 20 surface soils (directly outside of burrows) at 16 sites along Los Gatos Creek, and an additional 100 settled dust samples within the city of Coalinga. Our sampling effort was limited to accessible public land (parks, roadsides and waterways), and presents one of the most extensive such efforts among known studies (S1 Table). Coordinates were obtained using cellular phone GPS data cross-referenced with satellite imagery, and elevation data were estimated using elevatr version 0.4.2 (Hollister et al. 2022). All soils were collected as deeply as possible from within rodent burrows using steel hemispherical collectors attached to 30cm threaded rods, except for 20 soil samples from one site (Highway 33 West) along Los Gatos Creek that were collected from surface soil located adjacent to entrances of simultaneously sampled rodent burrows. For all soil samples, ≥25ml of soil was collected in conical 50ml polypropylene centrifuge tubes and mixed by repeatedly inverting the tubes. Sampling effort across sites is summarized in S2 Table.

We resampled a subset of locations to explore potential changes in *Coccidioides* presence over time (S3 Table). Previous research indicates that *Coccidioides* persists in rodent burrows or adjacent soils at the same location (Vargas-Gastelum et al. 2015, Wagner et al. 2023, Head et al. 2024). Along the Kern River, we sampled five sites (Kern River Parkway West End, University Place, Rio Kern Park, Yokuts Park, and Panorama Park West) at four time points over one year (April 2021, July 2021, October 2021 and January 2022). Where sites were sampled more than once, specific rodent burrows were not necessarily resampled. To investigate *Coccidioides* in soils unassociated with the Kern River (non-Kern River sites), we sampled four areas along California Highway 119, 10km south of the Kern River, at one time point each (Buena Vista Recreation Area, Buena Vista Valley, California State University Bakersfield (CSUB) Campus, and Sports Village). Along Los Gatos Creek, we sampled from the Clear Creek Management Area, through Coalinga, CA, to just north of Pleasant Valley State Prison near its confluence with Jacalitos Creek. Only one site was sampled more than once along Los Gatos Creek, Highway 33 West, in August 2020 and January 2022. In January 2022, we sampled east of California Highway 33 to the confluence of Los Gatos Creek with Warthan Creek and three sites near the confluence of Los Gatos Creek with Jacalitos Creek. In April 2024, we sampled ten sites along Los Gatos Creek Road along an elevational gradient from the Clear Creek Management Area in the Diablo Range, downhill to Coalinga, CA. In addition to soil samples, settled dust was sampled from 100 urban surfaces protected from precipitation by architectural features within Coalinga, CA, in August 2020. Settled dust was sampled by swabbing horizontal public surfaces using sterile, DNA-free cotton swabs (Puritan, Guilford, ME, USA). DNA was extracted from 0.25g of soil from each collection tube, and from swab tips cut from swab sticks. With both soil and swab tips, the samples were placed in MoBio Powersoil DNA kit buffer C1 (MoBio, Carlsbad, CA, USA) and disrupted in a FastPrep-24 5G bead beater (MP Biomedicals, Santa Ana, CA, USA), followed by DNA recovery using the MoBio Powersoil DNA kit as described (Wagner et al. 2022, 2023). DNA was diluted to 5ng µl^−1^ following measurement with the Qubit dsDNA HS Assay kit (Life Technologies Inc., Gaithersburg, MD, USA).

### Coccidioides detection

All samples were tested for *Coccidioides* using the CocciEnv qPCR assay (Bowers et al. 2019) in quadruplicate reactions, using nuclease-free water as a negative control and *C. posadasii* strain Silveira DNA as a positive control (provided by the lab of Anita Sil at the University of California San Francisco). The CocciENV assay builds upon the previously developed CocciDx assay (Litvintseva et al. 2015, Saubolle et al. 2018) by expanding the primer set targeting a unique repeating transposon sequence in the *Coccidioides* genome (NCBI BioProject PRJNA46299) and having been comprehensively validated using environmental soil samples (Bowers et al. 2019). Template DNA, 2μl (50ng·µl^−1^), was added to 10μl of TaqMan Environmental Master Mix2.0 (Applied Biosystems, Waltham, MA, USA), 2μl CocciEnv primer mix as described in Bowers et al. (2019), and 6μl nuclease-free H_2_O for a total of 20μl per reaction. The qPCR assay was performed on 96-well plates on the Stratagene Mx3000P platform (Agilent Technologies, Santa Clara, CA, USA) with the following cycling conditions: 1 cycle at 95°C for 10 minutes, 40 cycles at 95°C for 15 seconds each and 1 cycle at 60°C for 1 minute. Reactions with positive detections were identified at cycle threshold (CT) < 40, a minimum relative fluorescence unit value of 1000, and logarithmic amplification as described (Wagner et al. 2022, 2023). We were confident that an individual sample contained *Coccidioides* if 3 or more of its wells scored positive in the qPCR assay (Wagner et al. 2022, 2023). This threshold was chosen to balance sensitivity and specificity to minimize false positives while reliably detecting low-level presence. We also report the total number of positive wells for each sample (Fig 1, 2) (S2 Table, S3 Table). Settled dust recovered by swabbing provided a much smaller amount of sample compared to sampled soil, making it difficult to equate a failure to detect *Coccidioides* in settled dust with a failure to detect it in soil (Wagner et al. 2023). For this reason, following the protocol we established in Wagner et al. (2023), for settled dust samples having ≥1 positive well, multiple DNA extractions were concentrated using a vacuum oven operating at 20kPa and 65°C for one hour or until complete volume evaporation. Following resuspension of the concentrated DNA in nuclease-free H_2_O, a second analysis using the CocciEnv assay in quadruplicate reactions was done, and our initial analyses were updated for any samples that now showed *Coccidioides* detection in ≥ 3 replicate wells. All initial processing of qPCR data and generation of amplification curves used Mxpro version 4.1 (Agilent Technologies, Santa Clara, CA, USA).

### Fungal Community Amplicon Sequencing

To characterize the fungal soil community with regard to *Coccidioides* detection, amplicon sequencing of the internal transcribed (ITS2) region of fungal rDNA was performed on DNA extracted from 68 rodent burrow soil samples, 28 from four Kern River sites and 40 from one Los Gatos Creek site (20 within and 20 just outside the entrance of rodent burrows), all of which were also tested for *Coccidioides* using the CocciENV qPCR assay. An effort was made to perform sequencing on an equal number of *Coccidioides* positive and negative soils, as well as across an equal number of soils from two timepoints (April 2021 and July 2021) for Kern River samples and from two burrow locations (burrow interior and burrow entrance) for Los Gatos Creek soils (S4 Table). As previously described (Gao et al. 2018, Wagner et al. 2022), the ITS2 region was PCR amplified from extracted DNA using the 5.8SFun (AACTTTYRRCAAYGGATCWCT) and ITS4Fun (AGCCTCCGCTTATTGATATGCTTAART) primers (Taylor et al. 2016), with the AccuStart II PCR SuperMix kit (Quantabio, Beverly, MA, USA), on the Gene Amplification PCR System (Bio-Rad Laboratories, Hercules, CA, USA). The reaction mixture consisted of 2μl of template DNA, 2.5μl of 50μM forward and reverse primer each, 12.5μl AccuStart II PCR SuperMix, 2.5μl of nuclease-free water, and 3μl BSA, with the following thermal cycling conditions: an initial step of 96°C for 2 minutes, followed by 35 cycles of 94°C for 30 seconds, 58°C for 40 seconds and 72°C for 2 minutes, with a final step of 72°C for 10 minutes. The PCR product was quantified using the Qubit dsDNA HS Assay kit (Life Technologies Inc., Gaithersburg, MD, USA) and then sent to the QB3 Vincent J. Coates Genomics Sequencing Laboratory (University of California, Berkeley, CA, USA), where the samples were assigned unique dual indices to prevent barcode tag-jumping (Zinger et al. 2019, Carøe and Bohmann 2020) and were sequenced using paired-end PE300 chemistry on the MiSeq platform (Illumina, Inc., CA, USA).

### Amplicon Sequencing Data Processing

All sequence data were processed using Qiime 2 version 2019.10.0 (Bolyen et al. 2019). Quality control of the sequencing runs was performed manually followed by denoising using DADA2 (Callahan et al. 2016). Paired-end reads were then joined, end bases with a quality score lower than 25 were trimmed, and unpaired reads were removed. Assignment of operational taxonomic units (OTUs) to binned amplicon sequence variants (ASVs) was done with a naïve Bayes classifier, trained with the UNITE developer (untrimmed) database Qiime release version 9.0 (29.11.2022) clustered at 99% similarity (Pedregosa et al. 2011, Bokulich et al. 2018, Bolyen et al. 2019, Abarenkov et al. 2022). Sequences not assigned to specific taxonomies in the UNITE database were excluded in all downstream analyses. All supporting sequence data and metadata have been deposited in the NCBI Sequence Read Archive with BioProject numbers PRJNA1099691 and PRJNA1099703. All code needed to processing sequencing data and to generate taxonomic tables is included as a supplementary file (Code S1).

### Soil Moisture Data

To explore how the presence of *Coccidioides* in soils may coincide with precipitation and soil water availability (Kolivras and Comrie 2003, Comrie 2005, Tamerius and Comrie 2011), satellite-derived soil moisture data were acquired. Using the NASA AppEEARS user interface version 3.3.1 (AppEEARS Team 2022) (https://appeears.earthdatacloud.nasa.gov), the soil moisture active-passive (SMAP) L3 radar/radiometer global EASE-grid soil moisture dataset version 3 was downloaded, which is gridded at a 9km resolution using the EASE-Grid 2.0 projection, at a daily resolution (Entekhabi et al. 2016). SMAP L3 version 3 applies active–passive downscaling, model-based enhancements for vegetated or complex terrain and pixel overlap during satellite passes, all of which can introduce spatial detail at effective resolutions below 9km, depending on location. Surface (0-5cm) and rootzone (0-100cm) soil moisture data (cm^3^ water / cm^3^ soil) were selected using the “point sample tool” in the AppEEARS user interface at coordinates, both continuously along Lost Gatos Creek and the Kern River (surface only), as well as corresponding to soil sampling locations (both surface and rootzone), and mean values for each sampling month were calculated across sites. Due to the limited number of sites, and geographic range of sites along Los Gatos Creek, comparisons of site-level soil moisture data were limited to Bakersfield area sites, encompassing locations both directly along the Kern River and non-Kern River sites.

## Statistical Analysis

All statistical analyses used R version 4.1.0 (R Core Team 2020), vegan version 2.5.7 (Oksanen et al. 2019) and lme4 1.1.33 (Bates et al. 2015). Two separate logistic regression models were employed to determine significant differences in *Coccidioides* detection between sites, sampling timepoints and across remotely sensed soil moisture levels along the Kern River and surrounding areas. In the first logistic regression model, differences in *Coccidioides* detection were investigated as a function of remotely sensed soil moisture across all sites along the Kern River and the surrounding area. Sampling timepoint and elevation were included as fixed effects and, given the uneven sampling distribution (and dominant nature of Panorama Park West and University Place), site was included as a random effect to account for site-to-site variability. Surface and rootzone soil moisture and elevation were normalized by transforming each predictor into z-scores due to the large range in absolute values for each. In the second logistic regression model, differences in *Coccidioides* detection were tested between samples collected at two sites, Panorama Park West and University Place, which were sampled at all four time periods, and which had a high number of positive *Coccidioides* samples relative to other sites along the Kern River. Both sampling timepoint and remotely sensed soil moisture were included as covariates and z-score normalization was not performed as values between these two sites were relatively similar.

Given differences in the location and timing of sampling, patterns in soil fungal community structure were analyzed separately for Kern River and Los Gatos Creek sites. Species-level taxonomic count data was Wisconsin double transformed (Bray and Curtis 1957, Legendre and Legendre 2012) following the initial removal of species with only a single count across all samples, and Bray-Curtis dissimilarity was calculated as a measure of community β-diversity. To determine the correlation between fungal community β-diversity and *Coccidioides* detection, we employed a permutational (1000 permutations) multivariate analysis of variance (PERMANOVA) test, as implemented in the “adonis” function (Anderson 2001). Kern River samples included covariates for sampling site and timepoint. As the entire subset of Los Gatos Creek samples that underwent amplicon sequencing was collected from the same site, on the same day, only the specific sampling location (burrow interior versus burrow entrance) was included as a covariate. At Los Gatos Creek, soil moisture was excluded from PERMANOVA analysis due to the limited spatial extent of the subset of samples that was sequenced. We investigated possible co-occurrences between *Coccidioides* and other fungal taxa, extending our previous investigation of rodent burrow soils along California highway 33 (Wagner et al. 2023). An indicator species analysis was employed on untransformed species data using indicspecies version 1.7.12 (Dufrêne and Legendre 1997, De Cáceres et al. 2010) and significant results (p < 0.05, 1000 permutations) with ≥ 100 reads were cross-referenced using BLASTN (https://blast.ncbi.nlm.nih.gov) and Mycobank (https://www.mycobank.org) to determine likely alternative species identities (> 96% identity).

Figures were made with ggplot2 version 3.3.5 (Kahle and Wickham 2013), Complexheatmap version 2.8.0 (Gu et al. 2016), ggmap version 3.0.0 (Kahle and Wickham 2013) and osmdata version 0.1.8 (Padgham et al. 2017). Map data (lines) were acquired from OpenStreetMap (www.openstreetmap.org) under the Open Data Commons Open Database License (OdbL). Map tiles are by Stamen Design (www.stamen.com) under Creative Commons By Attribution (CC BY 4.0). California shapefile is public domain data by Natural Earth (www.naturalearthdata.com). Satellite imagery is public domain data provided by the United States Department of Agriculture (USDA) National Agriculture Imagery Program (NAIP) (https://naip-usdaonline.hub.arcgis.com). All non-sequencing data used in this study and all code needed to replicate statistical analyses are included as supplementary files (Data S1, S2; Code S2).

## Acknowledgements

This work was supported by National Institutes of Health, National Institute of Allergy and Infectious Disease grant R01AI148336, University of California Office of the President grant VFR-19-633952, and Department of Energy grant DE-SC0014081. We would like to thank Anita Sil’s lab for providing positive control *Coccidioides* DNA. The authors have no conflict of interest.

## Supporting Information Captions

**S1 Fig.** Map of urban surfaces sampled in Coalinga, CA in August 2020 and tested using the CocciENV qPCR assay. n = 100. Point colors correspond to positive replicate wells. Map data (lines) were acquired from OpenStreetMap (www.openstreetmap.org) under the Open Data Commons Open Database License (OdbL) (https://www.openstreetmap.org/copyright). Map tiles are by Stamen Design (www.stamen.com) under Creative Commons By Attribution (CC BY 4.0) (https://creativecommons.org/licenses/by/4.0/).

**S2 Fig.** Extended map of Los Gatos Creek rodent burrow soil sampling including April 2024 sites. All samples were tested using the CocciENV qPCR assay. Elevation profile (meters above sea level) and qualitative categorization of sample locations (a). Map of sample coordinates (b). Inset shows map extent. Landscape classification derived from the California Farmland Mapping and Monitoring Program 2018. Point colors correspond to positive replicate wells. Map data (lines) were acquired from OpenStreetMap (www.openstreetmap.org) under the Open Data Commons Open Database License (OdbL) (https://www.openstreetmap.org/copyright). Map tiles are by Stamen Design (www.stamen.com) under Creative Commons By Attribution (CC BY 4.0) (https://creativecommons.org/licenses/by/4.0/). California shapefile is public domain data by Natural Earth (www.naturalearthdata.com).

**S3 Fig.** The mean proportional abundance of the 30 most common fungal genera from sites along the Kern River as a function of site and sampling timepoint. (−) = *Coccidioides* negative samples. (+) = *Coccidioides* positive samples. n = 28.

**S4 Fig.** The mean proportional abundance of the 30 most common fungal genera from sites along Los Gatos Creek as a function of sampling location (rodent burrow entrance vs interior). (−) = *Coccidioides* negative samples. (+) = *Coccidioides* positive samples. n = 40.

**S1 Table.** Sampling enumeration for the current study (**bold text**) and all other studies known to the authors where environmental *Coccidioides* was detected in soil, air, or both. The location and the method of detection are shown here. Total studies (including the current study) = 42. (+) = positive samples.

**S2 Table.** *Coccidioides* detection in soils and settled dust using the CocciEnv qPCR assay as a function of site and detection level. Numbers in parentheses indicate the minimum proportion of replicate wells with CT < 40 and logarithmic amplification. Only samples detected in ≥ 3 replicate wells were considered positive detections. <u>*</u> = Bakersfield area / non-Kern River site

**S3 Table.** *Coccidioides* detection using the CocciEnv qPCR assay from sites with >1 timepoint as a function of site, date and detection level. Numbers in parentheses indicate the minimum proportion of replicate wells with CT < 40 and logarithmic amplification. * = Bakersfield area / non-Kern River site

**S4 Table.** Enumeration of subset of soil samples selected for ITS2 amplicon sequencing. n = 35 negative samples; 33 positive samples.

**S5 Table.** Generalized Linear Mixed Model (GLMM) coefficient table showing *Coccidioides* detection as a function of elevation, sampling month and soil moisture data (fixed effects), with site included as a random effect, across all Kern River sites. n = 830.

**S6 Table.** Modified generalized Linear Mixed Model (GLMM) coefficient table, excluding samples collected above 210 meters showing *Coccidioides* detection as a function of elevation, sampling month and soil moisture data (fixed effects), with site included as a random effect, across all Kern River sites (below 210 meters). n = 760.

**S7 Table.** Logistic regression coefficient table showing *Coccidioides* detection as a function of sampling site, sampling month and soil moisture data between Kern River sites at Panorama Park West and University Place. n = 202.

**S8 Table.** PERMANOVA coefficient table showing the association between the soil fungal community and *Coccidioides* detection (Presence), as a function of sampling site and sampling date, from rodent burrows soils collected from along the Kern River. *Coccidioides* detection determined using the CocciEnv qPCR assay. Fungal community structure modeled as species-level Bray-Curtis dissimilarities. Permutations = 1000. n = 28.

**S9 Table.** Pairwise fungal community differences (PERMANOVA) between Kern River sites using the ‘pairwiseadonis2’ function. Number of samples collected at each site shown in parentheses. Permutations = 1000. n = 28.

**S10 Table.** PERMANOVA coefficient table showing the association between the soil fungal community, *Coccidioides* detection (Presence) and sampling location (burrow interior vs entrance) from rodent burrow soils collected in August 2020 from the Los Gatos Creek “Hwy33 West” site. *Coccidioides* detection determined using the CocciEnv qPCR assay. Fungal community structure modeled as species-level Bray-Curtis dissimilarities. Permutations = 1000. n = 40.

**S11 Table.** Percent abundance of the 30 most abundant fungal genera at Kern River and Los Gatos Creek Sites. Values are means between replicates. Underlined genera appear uniquely in the top 30 genera at each location.

**S12 Table:** Indicator species for *Coccidioides* positive and *Coccidioides* negative soil samples collected from within Kern River rodent burrows. Significance was calculated in indicspecies version 1.7.12 with 1000 permutations. Sequences were cross-referenced via NCBI Nucleotide Blast (https://blast.ncbi.nlm.nih.gov/) for each prospective indicator species. • = equally likely alternative species (in subsection). IndVal and p-value apply to all alternative species.

**S13 Table:** Indicator species for *Coccidioides* positive soil samples collected from within Los Gato Creek rodent burrows. Significance was calculated in indicspecies version 1.7.12 with 1000 permutations. Sequences were cross-referenced via NCBI Nucleotide Blast (https://blast.ncbi.nlm.nih.gov/) for each prospective indicator species. • = equally likely alternative species (in subsection). IndVal and p-value apply to all alternative species.

**S14 Table:** Indicator species for *Coccidioides* negative soil samples collected from within Los Gatos Creek rodent burrows. Significance was calculated in indicspecies version 1.7.12 with 1000 permutations. Sequences were cross-referenced via NCBI Nucleotide Blast (https://blast.ncbi.nlm.nih.gov/) for each prospective indicator species. • = equally likely alternative species (in subsection). IndVal and p-value apply to all alternative species.

**S15 Table:** Indicator species for *Coccidioides* samples (positive and negative) collected from surface soils outside Los Gatos Creek rodent burrows. Significance was calculated in indicspecies version 1.7.12 with 1000 permutations. Sequences were cross-referenced via NCBI Nucleotide Blast (https://blast.ncbi.nlm.nih.gov/) for each prospective indicator species. • = equally likely alternative species (in subsection). IndVal and p-value apply to all alternative species.

**S1_data_qpcr_meta:** All qpcr data and all metadata.

**S2_data_soil_moisture:** All soil moisture data.

**S1_code_sequence_processing:** All code used to process raw sequencing data.

**S2_code_analysis:** All code used for statistical analyses.

