## Supplementary material for "Urban soils along the Kern River and Los Gatos Creek are hotspots for *Coccidioides* in the San Joaquin Valley of California": S1 Fig

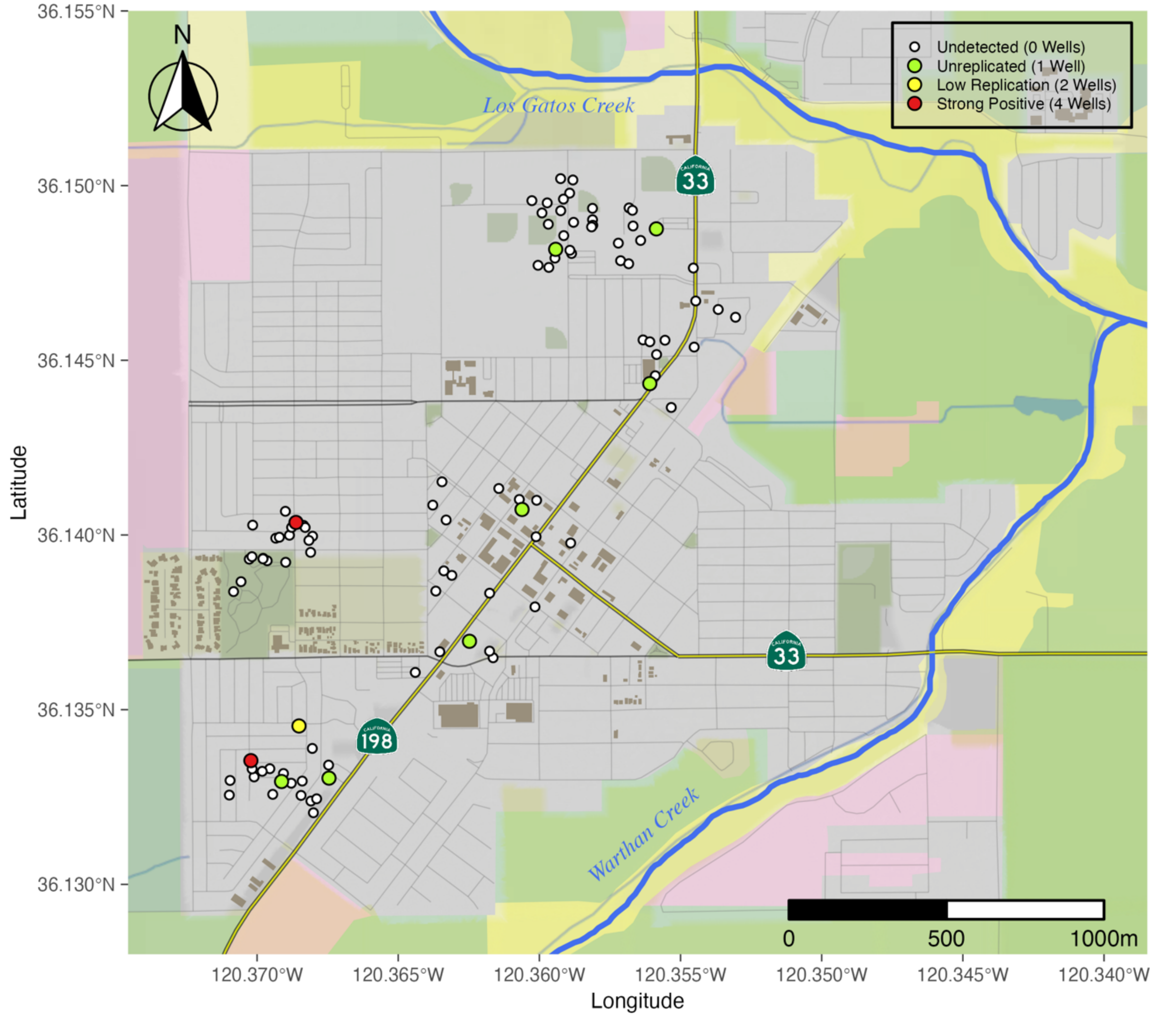


**S1 Fig.** Map of urban surfaces sampled in Coalinga, CA in August 2020 and tested using the CocciENV qPCR assay. n = 100. Point colors correspond to positive replicate wells. Map data (lines) were acquired from OpenStreetMap (www.openstreetmap.org) under the Open Data Commons Open Database License (OdbL) (https://www.openstreetmap.org/copyright). Map tiles are by Stamen Design (www.stamen.com) under Creative Commons By Attribution (CC BY 4.0) (https://creativecommons.org/licenses/by/4.0/).
