## Supplementary material for "Urban soils along the Kern River and Los Gatos Creek are hotspots for *Coccidioides* in the San Joaquin Valley of California": S1 Table

**S1 Table.** Sampling enumeration for the current study (**bold text**) and all other studies known to the authors where environmental *Coccidioides* was detected in soil, air, or both. The location and the method of detection are shown here. Total studies (including the current study) = 42. (+) = positive samples.

| | Collection Location | Detection Method | Total | (+) Total | Burrow | (+) Burrow | non-Burrow | (+) non-Burrow | Airborne | (+) Airborne |
|---|---|---|---|---|---|---|---|---|---|---|
| **Current Study** | **California** | **CocciENV qPCR** | **1178** | **117** | **1058** | **107** | **20** | **10** | **100** | **2** |
| Detection of Airborne Coccidioides Spores Using Lightweight Portable Air Samplers Affixed to Uncrewed Aircraft Systems in California's Central Valley (Radosevich et. al 2025) | California | CocciENV qPCR | 209 | 47 | 84 | 27 | 84 | 18 | 41 | 2 |
| Understanding the exposure risk of aerosolized Coccidioides in a Valley fever endemic metropolis (Porter et. al 2024) | Arizona | CocciENV qPCR | 5243 | 375 | 0 | 0 | 0 | 0 | 5243 | 375 |
| Small mammals and their burrows shape the distribution of Coccidioides in soils: a long-term ecological experiment." (Head et. al 2024) | California | CocciENV qPCR | 1988 | 322 | 1195 | 306 | 793 | 16 | 0 | 0 |
| Análisis de la distribución actual de Coccidioides spp. en muestras de suelo de Baja California, México (Mota 2024) | Mexico | Nested PCR, ddPCR | 76 | 5-30 | 76 | 5-30 | 0 | 0 | 0 | 0 |
| Coccidioides undetected in soils from agricultural land and uncorrelated with time or the greater soil fungal community on undeveloped land (Wagner et. al 2023) | California | CocciENV qPCR | 710 | 90 | 238 | 89 | 472 | 0 | 265 | 1 |
| Risk of Exposure to Coccidioides spp. in the Temblor Special Recreation Management Area (SRMA), Kern County, CA (Lauer et. al 2023) | California | Nested PCR | 114 | 36 | 15 | 5 | 99 | 31 | 0 | 0 |
| Locking Down the Dust: Exploring the Association Between Coccidioides posadasii and Biological Soil Crusts (Ramsey 2023) | Arizona | CocciDX qPCR | 80 | 2 | 20 | 2 | 60 | 0 | 0 | 0 |
| Coccidioidomycosis in Northern Arizona: an Investigation of the Host, Pathogen, and Environment Using a Disease Triangle Approach (Mead et. al 2022) | Arizona | CocciDX/ ENV qPCR | 171 | 6-26 | 171 | 6-26 | 0 | 0 | 0 | 0 |
| The detection of Coccidioides from ambient air in Phoenix, Arizona: Evidence of uneven distribution and seasonality (Gade et. al 2020) | Arizona | Single Tube Nested qPCR | 1009 | 0 | 0 | 0 | 0 | 0 | 1009 | 96-135 |
| Valley Fever: Environmental Risk Factors and Exposure Pathways Deduced from Field Measurements in California (Lauer et. al 2020) | California | Nested PCR | 389 | 122 | 0 | 0 | 389 | 122 | 0 | 0 |
| Investigating the Role of Animal Burrows on the Ecology and Distribution of Coccidioides spp. in Arizona Soils (Kollath et. al 2020) | Arizona | CocciDX/ ENV qPCR | 465 | 105 | 385 | 95 | 77 | 10 | 0 | 0 |
| Earthquake-ridden area in USA contains Coccidioides, the Valley fever pathogen (Lauer et. al 2020) | California | Nested PCR | 14 | 1 | 3 | 1 | 11 | 0 | 0 | 0 |
| Direct detection of Coccidioides from Arizona soils using CocciENV, a highly sensitive and specific real-time PCR assay (Bowers et. al 2019) | Arizona | CocciENV qPCR | 76 | 4 | 48 | 4 | 30 | 0 | 0 | 0 |
| Detection of Coccidioides posadasii from xerophytic environments in Venezuela reveals risk of naturally acquired coccidioidomycosis infections (Alvarado 2018) | Venezuela | GeneSTAT.MDx qPCR | 15 | 3-15 | - | - | - | - | 0 | 0 |
| Large-scale land development, fugitive dust, and increased coccidioidomycosis incidence in the Antelope Valley of California, 1999—2014 (Colson et. al 2017) | California | Nested PCR | 43 | 17 | - | - | - | - | 0 | 0 |
| Molecular detection of airborne Coccidioides in Tucson, Arizona (Chow et. al 2016) | Arizona | Single Tube Nested qPCR | 59 | 10 | - | 2 | - | 8 | 25 | 3 |
| Impact of seasonal changes on fungal diversity of a semi-arid ecosystem revealed by 454 pyrosequencing (Vargas-Gastélum et. al 2015) | Mexico | Amplicon Sequencing | 40 | 11 | 20 | 9 | 20 | 2 | 0 | 0 |
| Valley Fever: Finding New Places for an Old Disease: Coccidioides immitis Found in Washington State Soil Associated With Recent Human Infection (Litvintseva 2015) | Washington | CocciDX qPCR, Culture | 25 | 16 | - | - | - | - | 0 | 0 |
| Demonstration of Coccidioides immitis and Coccidioides posadasii DNA in soil samples collected from Dinosaur National Monument, Utah (Johnson et. al 2014) | Utah | Nested PCR | 2 | 2 | 1 | 1 | 1 | 1 | 0 | 0 |
| Combining Forces - The Use of Landsat TM Satellite Imagery, Soil Parameter Information, and Multiplex PCR to Detect | California | Multiplex PCR | ~192 | 12-144 | - | - | - | - | - | - |

| | | | | | | | | | | |
|---|---|---|---|---|---|---|---|---|---|---|
| Coccidioides immitis Growth Sites in Kern County, California (Lauer et. al 2014) | | | | | | | | | | |
| Coccidioides immitis identified in soil outside of its known range - Washington, 2013 (Marsden-Haug et. al 2013) | Washington | CocciDX qPCR | 22 | 6 | 0 | 0 | 0 | 0 | 0 | 0 |
| Molecular detection of Coccidioides spp. from environmental samples in Baja California: linking Valley Fever to soil and climate conditions (Baptista-Rosas et. al 2012) | Mexico | Nested PCR | 90 | 32 | 78 | 29 | 12 | 3 | 0 | 0 |
| Detection and phylogenetic analysis of Coccidioides posadasii in Arizona soil samples (Barker et. al 2012) | Arizona | PCR, Mouse, Culture | 700 | 11 | - | 5 | - | 6 | 0 | 0 |
| Detection of Coccidioides immitis in Kern County, California, by multiplex PCR (Lauer et. al 2011) | California | PCR | 546 | 31 | 0 | 0 | 546 | 31 | 0 | 0 |
| Molecular identification of Coccidioides spp. in soil samples from Brazil (Macêdo et. al 2011) | Brazil | Nested PCR, Mouse Injection | 24 | 6 | 24 | 6 | 0 | 0 | 0 | 0 |
| Genomic and Population Analyses of the Mating Type Loci in Coccidioides Species Reveal Evidence for Sexual Reproduction and Gene Acquisition (Mandel et. al 2007) | Arizona | Mouse Injection | 11 | 11 | - | - | - | - | - | - |
| Phenotypic characterization and ecological features of Coccidioides spp. from Northeast Brazil (Cordeiro et. al 2006) | Brazil | Culture | 14 | 3 | 14 | 3 | 0 | 0 | 0 | 0 |
| Development of a Quantitative TaqMan™-PCR Assay and Feasibility of Atmospheric Collection for Coccidioides immitis for Ecological Studies (Daniels et. al 2002) | California | PCR | 0 | 0 | 0 | 0 | 0 | 0 | 12 | 4 |
| Outbreak of Coccidioidomycosis in Washington State Residents Returning from Mexico (Cairns et. al 2000) | Mexico | Mouse Injection | 3 | 3 | - | - | - | - | - | - |
| Soil isolation and molecular identification of Coccidioides immitis (Greene et. al 2000) | California | PCR | 720 | 4 | 0 | 0 | 720 | 4 | 0 | 0 |
| Soil Ecology of Coccidioides immitis at Amerindian Middens in California (Lacy and Swatek 1974) | California | Culture | 325 | 32 | 0 | 0 | 325 | 32 | 0 | 0 |
| Some fungi isolated with Coccidioides immitis from soils of endemic areas in California (Orr 1968) | California | Culture | 22 | 22 | 0 | 0 | 22 | 22 | 0 | 0 |
| Recovery of Coccidioides immitis from the air (Ajello et. al 1965) | Arizona | Mouse Injection, Culture | 0 | 0 | 0 | 0 | 0 | 0 | 128 | 2 |
| Observations on Coccidioides immitis found growing naturally in soil. (Maddy et. al 1965) | Arizona | Culture, Injection | 198 | 30 | 0 | 0 | 198 | 30 | 0 | 0 |
| Isolation of Coccidioides immitis from Soil (Levine et. al 1964) | California | Culture, Mouse Injection | 37 | 4 | - | - | - | - | 0 | 0 |
| Growth Patterns of Coccidioides immitis (Elconin 1957) | California | Culture, Mouse | 428 | 31 | 115 | 18 | 313 | 13 | 0 | 0 |
| Ecological studies of Coccidioides immitis (Plunkett and Swatek 1957) | California | Culture | 80 | 0 | 80 | 0 | 0 | 0 | 0 | 0 |
| Coccidioides immitis in the soil of the southern San Joaquin Valley. (Egeberg and Ely 1954) | California | Culture, Injection | 500 | 35 | 177 | 24 | 323 | 11 | 0 | 0 |
| Isolation of Coccidioides from soil and rodents (Emmons 1942) | Arizona | Isolation | 150 | 5 | - | - | - | - | 0 | 0 |
| An Epidemic Of Coccidioidal Infection (Coccidioidomycosis) (Davis et. al 1942) | California | Isolation | 1 | 1 | 1 | 0 | 0 | 0 | 0 | 0 |
| Isolation of Coccidioides Immitis (Stiles) from the Soil (Stewart and Meyer 1932) | California | Culture, Guinea Pig | >1 | >1 | - | - | - | - | 0 | 0 |
