## Supplementary material for "Urban soils along the Kern River and Los Gatos Creek are hotspots for *Coccidioides* in the San Joaquin Valley of California": S2 Fig

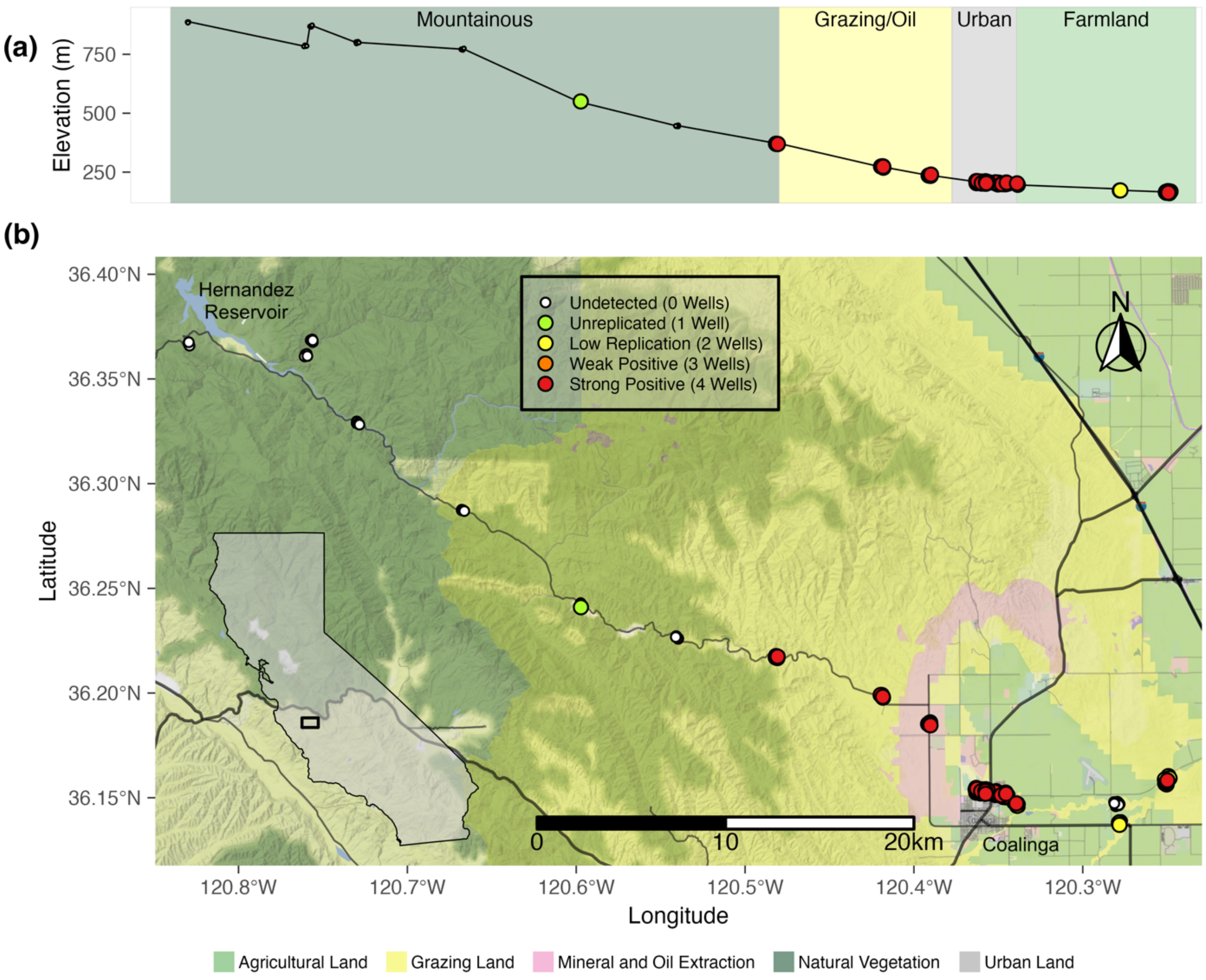


**S2 Fig.** Extended map of Los Gatos Creek rodent burrow soil sampling including April 2024 sites. All samples were tested using the CocciENV qPCR assay. Elevation profile (meters above sea level) and qualitative categorization of sample locations (a). Map of sample coordinates (b). Inset shows map extent. Landscape classification derived from the California Farmland Mapping and Monitoring Program 2018. Point colors correspond to positive replicate wells. Map data (lines) were acquired from OpenStreetMap (www.openstreetmap.org) under the Open Data Commons Open Database License (OdbL) (https://www.openstreetmap.org/copyright). Map tiles are by Stamen Design (www.stamen.com) under Creative Commons By Attribution (CC BY 4.0) (https://creativecommons.org/licenses/by/4.0/). California shapefile is public domain data by Natural Earth (www.naturalearthdata.com).
