## Supplementary material for "Urban soils along the Kern River and Los Gatos Creek are hotspots for *Coccidioides* in the San Joaquin Valley of California": S2 Table

**S2 Table.** *Coccidioides* detection in soils and settled dust using the CocciEnv qPCR assay as a function of site and detection level. Numbers in parentheses indicate the minimum proportion of replicate wells with CT < 40 and logarithmic amplification. Only samples detected in ≥ 3 replicate wells were considered positive detections. * = Bakersfield area / non-Kern River site

| | Undetected (0/4) | Unreplicated (1/4) | Low Replication (2/4) | Weak Positive (3/4) | Strong Positive (4/4) | Total |
|---|---|---|---|---|---|---|
| **Kern River (soils)** | | | | | | |
| 1. Buena Vista Valley * (non-Kern River site) | 13 | 2 | 0 | 0 | 0 | 15 |
| 2. California Aqueduct * (non-Kern River site) | 8 | 4 | 0 | 0 | 0 | 12 |
| 3. Buena Vista Recreation Area * (non-Kern River | 45 | 1 | 0 | 0 | 0 | 46 |
| 4. Kern River Mouth | 1 | 1 | 0 | 0 | 0 | 2 |
| 5. Kern River Parkway West End | 59 | 1 | 1 | 1 | 1 | 63 |
| 6. Highgate Park | 3 | 0 | 0 | 0 | 0 | 3 |
| 7. River Walk | 4 | 2 | 0 | 0 | 0 | 6 |
| 8. CSUB West * (non-Kern River site) | 38 | 0 | 0 | 0 | 0 | 38 |
| 9. CSUB North * (non-Kern River site) | 7 | 0 | 0 | 0 | 0 | 7 |
| 10. University Place | 68 | 7 | 6 | 4 | 21 | 106 |
| 11. Sports Village West * (non-Kern River site) | 20 | 0 | 0 | 0 | 0 | 20 |
| 12. Sports Village East * (non-Kern River site) | 65 | 2 | 0 | 0 | 0 | 67 |
| 13. Rio Kern Park | 55 | 5 | 0 | 2 | 0 | 62 |
| 14. Truxtun Park | 20 | 0 | 0 | 0 | 0 | 20 |
| 15. Yokuts Park | 60 | 1 | 2 | 0 | 0 | 63 |
| 16. Beach Park | 26 | 0 | 0 | 0 | 0 | 26 |
| 17. Upland Park | 3 | 0 | 0 | 0 | 0 | 3 |
| 18. Panorama Vista Preserve | 3 | 0 | 0 | 0 | 0 | 3 |
| 19. Panorama Park West | 75 | 0 | 1 | 0 | 20 | 96 |
| 20. Greenlawn Cemetery | 6 | 0 | 0 | 0 | 0 | 6 |
| 21. Panorama Park East | 17 | 1 | 0 | 0 | 2 | 20 |
| 22. Kern River Parkway East Bridge | 2 | 1 | 0 | 0 | 0 | 3 |
| 23. Kern River Parkway East End | 21 | 0 | 0 | 1 | 0 | 22 |
| 24. Gordons Ferry | 1 | 2 | 0 | 0 | 0 | 3 |
| 25. Round Mountain Road | 3 | 0 | 0 | 0 | 0 | 3 |
| 26. Hart Park West | 3 | 0 | 0 | 0 | 0 | 3 |
| 27. Hart Park East | 3 | 0 | 0 | 0 | 0 | 3 |
| 28. Kern County Soccer Park | 29 | 0 | 0 | 0 | 0 | 29 |
| 29. Kern River Golf Course | 4 | 0 | 0 | 0 | 0 | 4 |
| 30. Rancheria Road | 6 | 0 | 0 | 0 | 0 | 6 |
| 31. Upper Richbar | 6 | 0 | 0 | 0 | 0 | 6 |
| 32. Kern Canyon Road West End | 6 | 0 | 0 | 0 | 0 | 6 |
| 33. Kern Canyon Road 4km | 6 | 0 | 0 | 0 | 0 | 6 |
| 34. Kern Canyon Road 11km | 6 | 0 | 0 | 0 | 0 | 6 |
| 35. Remington Ridge Trailhead | 7 | 0 | 0 | 0 | 0 | 7 |
| 36. Hobo Campground | 3 | 0 | 0 | 0 | 0 | 3 |
| 37. Sandy Flat Campground | 7 | 0 | 0 | 0 | 0 | 7 |
| 38. Tank Park | 6 | 0 | 0 | 0 | 0 | 6 |
| 39. Isabella Hot Springs | 5 | 0 | 0 | 0 | 0 | 5 |
| 40. Auxiliary Dam Campground | 6 | 0 | 0 | 0 | 0 | 6 |
| 41. Old Isabella | 6 | 0 | 0 | 0 | 0 | 6 |
| 42. South Fork Rec Campground | 6 | 0 | 0 | 0 | 0 | 6 |
| | **738** | **30** | **10** | **8** | **44** | **830** |
| **Los Gatos Creek (soils)** | | | | | | |
| W01. Laguna Mountain Campground | 3 | 0 | 0 | 0 | 0 | 3 |
| W02. Oak Flat Campground | 4 | 0 | 0 | 0 | 0 | 4 |
| W03. Jade Mill Campground | 6 | 0 | 0 | 0 | 0 | 6 |
| W04. Mile 4.3 | 5 | 0 | 0 | 0 | 0 | 5 |
| W05. Mile 9.4 | 5 | 0 | 0 | 0 | 0 | 5 |
| W06. Benitoite Mine | 4 | 1 | 0 | 0 | 0 | 5 |
| W07. Mile 19.5 | 5 | 0 | 0 | 0 | 0 | 5 |
| W08. Mile 24 | 1 | 0 | 0 | 0 | 4 | 5 |
| W09. Mile 28.6 | 0 | 0 | 0 | 0 | 5 | 5 |
| W10. Mile 31.1 | 0 | 0 | 0 | 3 | 2 | 5 |
| 1A. Highway 33 West (burrows) | 34 | 3 | 6 | 4 | 13 | 60 |
| 1B. Highway 33 West (surface soils) | 8 | 0 | 2 | 4 | 6 | 20 |
| 2. Highway 33 East | 24 | 0 | 1 | 1 | 14 | 40 |
| 3. Warthan Creek | 18 | 0 | 0 | 1 | 1 | 20 |
| 4. South Calavaras | 20 | 0 | 0 | 0 | 0 | 20 |
| 5. Jacalitos Creek | 18 | 1 | 1 | 0 | 0 | 20 |
| 6. Phelps Avenue 10km | 11 | 2 | 2 | 0 | 5 | 20 |
| | **166** | **7** | **12** | **13** | **50** | **248** |
| **Coalinga (urban surfaces)** | | | | | | |
| 1. Coalinga | 90 | 7 | 1 | 0 | 2 | 100 |
| | **90** | **7** | **1** | **0** | **2** | **100** |
| **Total** | 994 | 44 | 23 | 21 | 96 | 1178 |
