## Supplementary material for "Urban soils along the Kern River and Los Gatos Creek are hotspots for *Coccidioides* in the San Joaquin Valley of California": S3 Fig

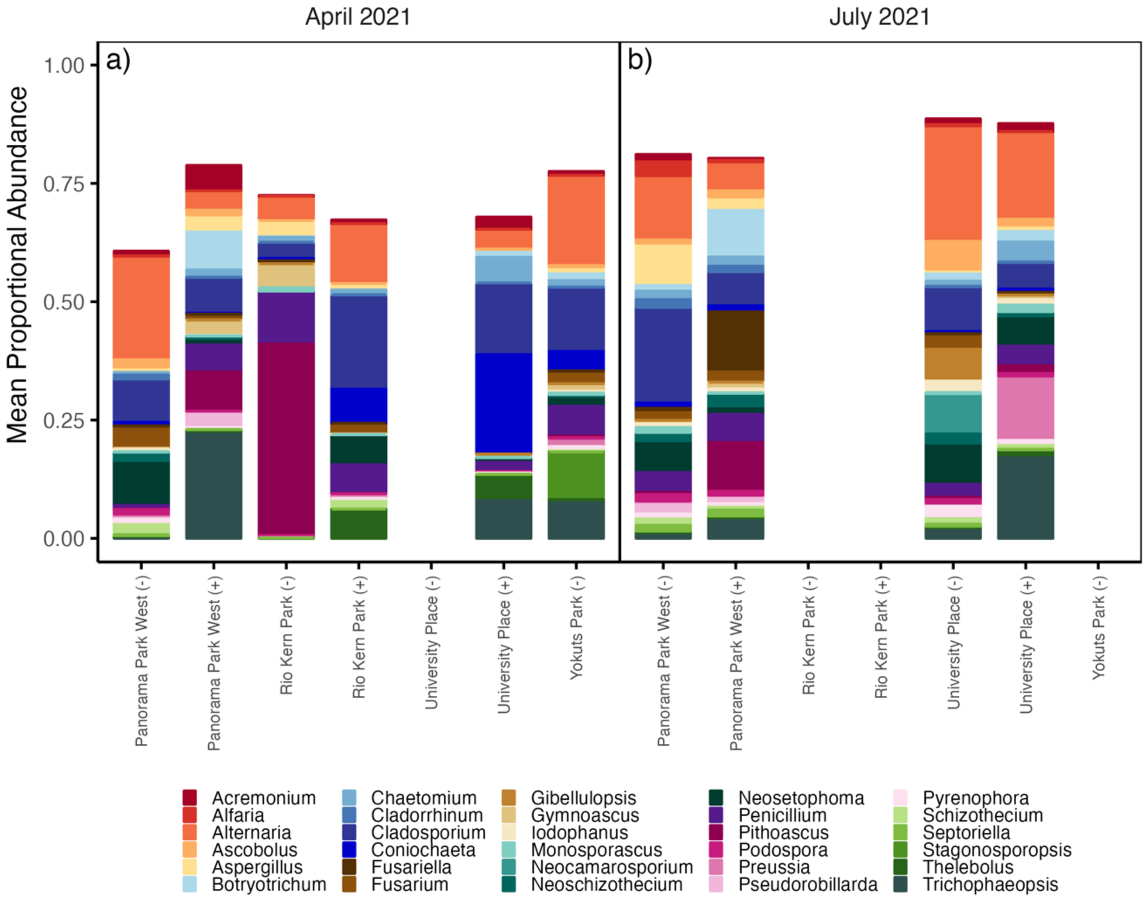


**S3 Fig.** The mean proportional abundance of the 30 most common fungal genera from sites along the Kern River as a function of site and sampling timepoint. (-) = *Coccidioides* negative samples. (+) = *Coccidioides* positive samples. n = 28.
