## Supplementary material for "Urban soils along the Kern River and Los Gatos Creek are hotspots for *Coccidioides* in the San Joaquin Valley of California": S3 Table

**S3 Table.** *Coccidioides* detection using the CocciEnv qPCR assay from sites with >1 timepoint as a function of site, date and detection level. Numbers in parentheses indicate the minimum proportion of replicate wells with CT < 40 and logarithmic amplification. * = Bakersfield area / non-Kern River site

| | | Undetected (0/4) | Unreplicated (1/4) | Low Replication (2/4) | Weak Positive (3/4) | Strong Positive (4/4) | Total |
|---|---|---|---|---|---|---|---|
| **Kern River (soils)** | | | | | | | |
| 3. Buena Vista Recreation Area * (non-Kern River site) | Apr 2021 | 6 | 0 | 0 | 0 | 0 | 6 |
| | Jan 2022 | 39 | 1 | 0 | 0 | 0 | 40 |
| 5. Kern River Parkway West End | Apr 2021 | 2 | 1 | 0 | 0 | 0 | 3 |
| | Jul 2021 | 20 | 0 | 0 | 0 | 0 | 20 |
| | Oct 2021 | 20 | 0 | 0 | 0 | 0 | 20 |
| | Jan 2022 | 17 | 0 | 1 | 1 | 1 | 20 |
| 8. CSUB West * (non-Kern River site) | Jul 2021 | 20 | 0 | 0 | 0 | 0 | 20 |
| | Oct 2021 | 18 | 0 | 0 | 0 | 0 | 18 |
| 9. CSUB North * (non-Kern River site) | Jul 2021 | 4 | 0 | 0 | 0 | 0 | 4 |
| | Oct 2021 | 3 | 0 | 0 | 0 | 0 | 3 |
| 10. University Place | Apr 2021 | 1 | 4 | 0 | 1 | 0 | 6 |
| | Jul 2021 | 14 | 1 | 1 | 0 | 4 | 20 |
| | Oct 2021 | 16 | 0 | 0 | 0 | 4 | 20 |
| | Jan 2022 | 37 | 2 | 5 | 3 | 13 | 60 |
| 12. Sports Village East * (non-Kern River site) | Apr 2021 | 24 | 2 | 0 | 0 | 0 | 26 |
| | Jul 2021 | 21 | 0 | 0 | 0 | 0 | 21 |
| | Oct 2021 | 20 | 0 | 0 | 0 | 0 | 20 |
| 13. Rio Kern Park | Apr 2021 | 1 | 1 | 0 | 1 | 0 | 3 |
| | Jul 2021 | 19 | 1 | 0 | 0 | 0 | 20 |
| | Oct 2021 | 18 | 1 | 0 | 0 | 0 | 19 |
| | Jan 2022 | 17 | 2 | 0 | 1 | 0 | 20 |
| 15. Yokuts Park | Apr 2021 | 2 | 0 | 1 | 0 | 0 | 3 |
| | Jul 2021 | 19 | 1 | 0 | 0 | 0 | 20 |
| | Oct 2021 | 19 | 0 | 1 | 0 | 0 | 20 |
| | Jan 2022 | 20 | 0 | 0 | 0 | 0 | 20 |
| 16. Beach Park | Apr 2021 | 6 | 0 | 0 | 0 | 0 | 6 |
| | Jan 2022 | 20 | 0 | 0 | 0 | 0 | 20 |
| 19. Panorama Park West | Apr 2021 | 4 | 0 | 0 | 0 | 2 | 6 |
| | Jul 2021 | 15 | 0 | 1 | 0 | 4 | 20 |
| | Oct 2021 | 26 | 0 | 0 | 0 | 4 | 30 |
| | Jan 2022 | 30 | 0 | 0 | 0 | 10 | 40 |
| 28. Kern County Soccer Park | Apr 2021 | 9 | 0 | 0 | 0 | 0 | 9 |
| | Oct 2021 | 20 | 0 | 0 | 0 | 0 | 20 |
| | | **527** | **17** | **10** | **7** | **42** | **603** |
| **Los Gatos Creek (soils)** | | | | | | | |
| 1A. Highway 33 West (burrows) | Aug 2020 | 6 | 2 | 1 | 3 | 8 | 20 |
| | Jan 2022 | 28 | 1 | 5 | 1 | 5 | 40 |
| 1B. Highway 33 West (surface soils) | Aug 2020 | 8 | 0 | 2 | 4 | 6 | 20 |
| | | **42** | **3** | **8** | **8** | **19** | **80** |
| **Total** | | **569** | **20** | **18** | **15** | **61** | **683** |
