## Supplementary material for "Urban soils along the Kern River and Los Gatos Creek are hotspots for *Coccidioides* in the San Joaquin Valley of California": S4 Fig

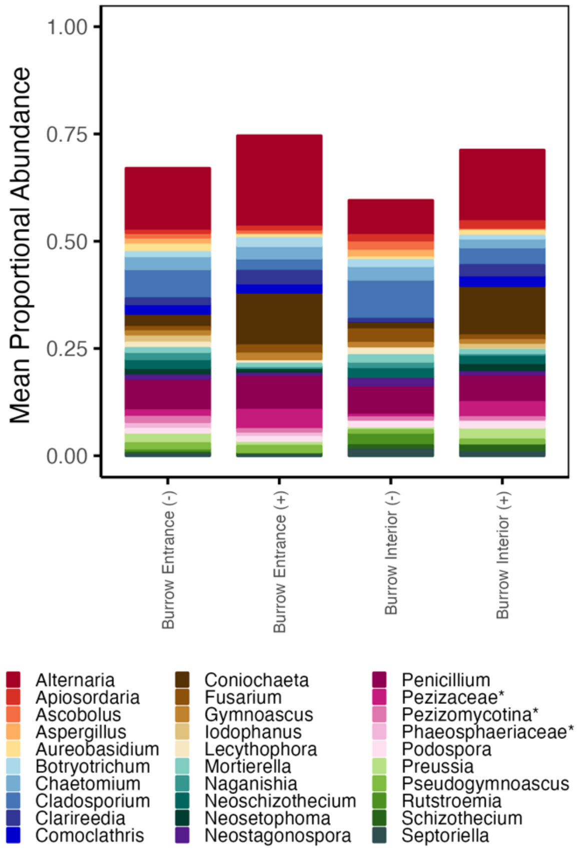


**S4 Fig.** The mean proportional abundance of the 30 most common fungal genera from sites along Los Gatos Creek as a function of sampling location (rodent burrow entrance vs interior). (-) = *Coccidioides* negative samples. (+) = *Coccidioides* positive samples. n = 40.
