## Supplementary material for "Urban soils along the Kern River and Los Gatos Creek are hotspots for *Coccidioides* in the San Joaquin Valley of California": S4 Table

**S4 Table.** Enumeration of subset of soil samples selected for ITS2 amplicon sequencing. n = 35 negative samples; 33 positive samples.

| Waterway | Burrow position | Timepoint | Site | *Coccidioides* | # of Samples |
|---|---|---|---|---|---|
| Los Gatos Creek | Burrow Entrance | Aug 2020 | Highway 33 West | Negative (-) | 10 |
| Los Gatos Creek | Burrow Interior | Aug 2020 | Highway 33 West | Negative (-) | 9 |
| Kern River | Burrow Interior | Apr 2021 | Panorama Park West | Negative (-) | 2 |
| Kern River | Burrow Interior | Jul 2021 | Panorama Park West | Negative (-) | 6 |
| Kern River | Burrow Interior | Apr 2021 | Rio Kern Park | Negative (-) | 1 |
| Kern River | Burrow Interior | Jul 2021 | University Place | Negative (-) | 4 |
| Kern River | Burrow Interior | Apr 2021 | Yokuts Park | Negative (-) | 3 |
| Los Gatos Creek | Burrow Entrance | Aug 2020 | Highway 33 West | Positive (+) | 10 |
| Los Gatos Creek | Burrow Interior | Aug 2020 | Highway 33 West | Positive (+) | 11 |
| Kern River | Burrow Interior | Apr 2021 | Panorama Park West | Positive (+) | 2 |
| Kern River | Burrow Interior | Jul 2021 | Panorama Park West | Positive (+) | 4 |
| Kern River | Burrow Interior | Apr 2021 | Rio Kern Park | Positive (+) | 1 |
| Kern River | Burrow Interior | Apr 2021 | University Place | Positive (+) | 1 |
| Kern River | Burrow Interior | Jul 2021 | University Place | Positive (+) | 4 |
| **Total** | | | | | **68** |
