## Supplementary material for "Urban soils along the Kern River and Los Gatos Creek are hotspots for *Coccidioides* in the San Joaquin Valley of California": S5 Table

**S5 Table.** Generalized Linear Mixed Model (GLMM) coefficient table showing *Coccidioides* detection as a function of elevation, sampling month and soil moisture data (fixed effects), with site included as a random effect, across all Kern River sites. n = 830.

| | Estimate | OR | CI 2.5% | CI 97.5% | SE | z-value | p-value | |
|---|---|---|---|---|---|---|---|---|
| Intercept | -6.926 | 9.82E-04 | 3.36E-05 | 0.03 | 1.721 | -4.024 | 0 | *** |
| Elevation | 4.463 | 86.75 | 0.62 | 1.21E+04 | 2.518 | 1.773 | 0.076 | . |
| Surface Soil Moisture (0-5cm) | -2.736 | 0.06 | 4.25E-03 | 0.99 | 1.39 | -1.968 | 0.049 | * |
| Rootzone Soil Moisture (0-100cm) | -4.786 | 8.35E-03 | 2.61E-05 | 2.67 | 2.943 | -1.626 | 0.104 | |
| 2021 July | -4.028 | 0.02 | 3.42E-04 | 0.93 | 2.017 | -1.996 | 0.046 | * |
| 2021 October | -3.817 | 0.02 | 5.89E-04 | 0.82 | 1.847 | -2.066 | 0.039 | * |
| 2022 January | 7.834 | 2.52E+03 | 1.40 | 4.54E+06 | 3.824 | 2.049 | 0.04 | * |

. = $p < 0.1$, * = $p < 0.05$, ** = $p < 0.01$, *** = $p \leq 0.001$, OR = Odds Ratio, CI = Confidence Interval, SE = Standard Error
