## Supplementary material for "Urban soils along the Kern River and Los Gatos Creek are hotspots for *Coccidioides* in the San Joaquin Valley of California": S6 Table

**S6 Table.** Modified generalized Linear Mixed Model (GLMM) coefficient table, excluding samples collected above 210 meters showing *Coccidioides* detection as a function of elevation, sampling month and soil moisture data (fixed effects), with site included as a random effect, across all Kern River sites (below 210 meters). n = 760.

| | Estimate | OR | CI 2.5% | CI 97.5% | SE | z-value | p-value | |
|---|---|---|---|---|---|---|---|---|
| Intercept | -13.844 | 4.96E-03 | 2.09E-04 | 0.12 | 4.204 | -3.293 | 0.001 | *** |
| Elevation | 0.048 | 3.70E+03 | 5.26 | 2.61E+06 | 0.02 | 2.456 | 0.014 | * |
| Surface Soil Moisture (0-5cm) | -2.272 | 0.10 | 7.16E-03 | 1.48 | 1.361 | -1.67 | 0.095 | . |
| Rootzone Soil Moisture (0-100cm) | -4.287 | 0.01 | 7.65E-05 | 2.47 | 2.649 | -1.619 | 0.106 | |
| 2021 July | -3.594 | 0.03 | 6.82E-04 | 1.11 | 1.887 | -1.905 | 0.057 | . |
| 2021 October | -3.343 | 0.04 | 1.21E-03 | 1.03 | 1.721 | -1.942 | 0.052 | . |
| 2022 January | 6.869 | 961.62 | 0.88 | 1.06E+06 | 3.573 | 1.923 | 0.055 | . |

. = $p < 0.1$, * = $p < 0.05$, ** = $p < 0.01$, *** = $p \leq 0.001$, OR = Odds Ratio, CI = Confidence Interval, SE = Standard Error
