## Supplementary material for "Urban soils along the Kern River and Los Gatos Creek are hotspots for *Coccidioides* in the San Joaquin Valley of California": S7 Table

**S7 Table.** Logistic regression coefficient table showing *Coccidioides* detection as a function of sampling site, sampling month and soil moisture data between Kern River sites at Panorama Park West and University Place. n = 202.

| | Estimate | OR | CI 2.5% | CI 97.5% | SE | z-value | p-value |
|---|---|---|---|---|---|---|---|
| Intercept | -4.878 | 0.01 | 5.35E-24 | 3.82E+18 | 24.209 | -0.201 | 0.84 |
| July 2021 | 0.899 | 2.46 | 0.00 | 4.88E+06 | 7.21 | 0.125 | 0.90 |
| October 2021 | 0.587 | 1.80 | 0.00 | 1.34E+06 | 6.673 | 0.088 | 0.93 |
| January 2022 | -2.055 | 0.13 | 1.91E-13 | 7.46E+10 | 13.647 | -0.151 | 0.88 |
| Site (Panorama Park West) | -1.105 | 0.33 | 0.00 | 1.99E+04 | 5.571 | -0.198 | 0.84 |
| Surface Soil Moisture (0-5cm) | 12.703 | 3.29E+05 | 5.62E-104 | 7.20E+115 | 127.011 | 0.1 | 0.92 |
| Rootzone Soil Moisture (0-100cm) | 44.459 | 2.03E+19 | 1.63E-193 | 1.08E+235 | 247.267 | 0.18 | 0.86 |

. = $p < 0.1$, * = $p < 0.05$, ** = $p < 0.01$, *** = $p \leq 0.001$, OR = Odds Ratio, CI = Confidence Interval, SE = Standard Error
