## Supplementary material for "Urban soils along the Kern River and Los Gatos Creek are hotspots for *Coccidioides* in the San Joaquin Valley of California": S8 Table

**S8 Table.** PERMANOVA coefficient table showing the association between the soil fungal community and *Coccidioides* detection (Presence), as a function of sampling site and sampling date, from rodent burrows soils collected from along the Kern River. *Coccidioides* detection determined using the CocciEnv qPCR assay. Fungal community structure modeled as species-level Bray-Curtis dissimilarities. Permutations = 1000. n = 28.

| | Degrees of Freedom | Sum of Squares | $r^2$ | Pseudo-F | p-value | |
|---|---|---|---|---|---|---|
| *Coccidioides* Presence | 1 | 0.327 | 0.03 | 1.067 | 0.395 | |
| Site | 3 | 2.145 | 0.22 | 2.335 | 0.002 | ** |
| Date | 1 | 0.394 | 0.04 | 1.286 | 0.234 | |
| *Coccidioides* Presence : Site | 2 | 0.83 | 0.09 | 1.355 | 0.145 | |
| *Coccidioides* Presence : Date | 1 | 0.217 | 0.02 | 0.71 | 0.702 | |
| Site : Date | 1 | 0.382 | 0.04 | 1.247 | 0.253 | |
| Residual | 18 | 5.512 | 0.56 | | | |
| **Total** | **27** | **9.807** | **1** | | | |

. = $p < 0.1$, * = $p < 0.05$, ** = $p < 0.01$, *** = $p \leq 0.001$
