## Supplementary material for "Urban soils along the Kern River and Los Gatos Creek are hotspots for *Coccidioides* in the San Joaquin Valley of California": S9 Table

**S9 Table.** Pairwise fungal community differences (PERMANOVA) between Kern River sites using the 'pairwiseadonis2' function. Number of samples collected at each site shown in parentheses. Permutations = 1000. n = 28.

| Contrast | Degrees of Freedom | Sum of Squares | $r^2$ | Pseudo-F | p-value | |
|---|---|---|---|---|---|---|
| Panorama Park West (14) : University Place (9) | 1 | 0.7 | 0.08 | 1.72 | 0.04 | * |
| Panorama Park West (14) : Rio Kern Park (2) | 1 | 0.35 | 0.06 | 0.82 | 0.71 | |
| Panorama Park West (14) : Yokuts Park (3) | 1 | 0.42 | 0.06 | 0.99 | 0.42 | |
| University Place (9) : Rio Kern Park (2) | 1 | 0.23 | 0.06 | 0.58 | 0.93 | |
| University Place (9) : Yokuts Park (3) | 1 | 0.43 | 0.10 | 1.09 | 0.32 | |
| Rio Kern Park (2) : Yokuts Park (3) | 1 | 0.42 | 0.25 | 0.98 | 0.40 | |

. = $p < 0.1$, * = $p < 0.05$, ** = $p < 0.01$, *** = $p \leq 0.001$
