## Supplementary material for "Urban soils along the Kern River and Los Gatos Creek are hotspots for *Coccidioides* in the San Joaquin Valley of California": S10 Table

**S10 Table.** PERMANOVA coefficient table showing the association between the soil fungal community, *Coccidioides* detection (Presence) and sampling location (burrow interior vs entrance) from rodent burrow soils collected in August 2020 from the Los Gatos Creek "Hwy33 West" site. *Coccidioides* detection determined using the CocciEnv qPCR assay. Fungal community structure modeled as species-level Bray-Curtis dissimilarities. Permutations = 1000. n = 40.

| | Degrees of Freedom | Sum of Squares | $r^2$ | Pseudo-F | p-value |
|---|---|---|---|---|---|
| *Coccidioides* Presence | 1 | 0.373 | 0.03 | 1.174 | 0.234 |
| Sampling Location | 1 | 0.318 | 0.03 | 1 | 0.381 |
| *Coccidioides* Presence : Sampling Location | 1 | 0.269 | 0.02 | 0.846 | 0.682 |
| Residual | 36 | 11.435 | 0.92 | | |
| **Total** | **39** | **12.394** | **1** | | |

. = $p < 0.1$, * = $p < 0.05$, ** = $p < 0.01$, *** = $p \leq 0.001$
