## Supplementary material for "Urban soils along the Kern River and Los Gatos Creek are hotspots for *Coccidioides* in the San Joaquin Valley of California": S11 Table

**S11 Table.** Percent abundance of the 30 most abundant fungal genera at Kern River and Los Gatos Creek Sites. Values are means between replicates. Underlined genera appear uniquely in the top 30 genera at each location.

| Kern River | | | Los Gatos Creek | | |
|---|---|---|---|---|---|
| | mean | sem | | mean | sem |
| Alternaria | 14.58 | 2.05 | Alternaria | 15.38 | 1.67 |
| Cladosporium | 11.51 | 1.61 | Coniochaeta | 7.6 | 1.81 |
| Trichophaeopsis | 6.53 | 2.6 | Penicillium | 7.35 | 1.61 |
| Neosetophoma | 5.09 | 1.31 | Cladosporium | 5.78 | 0.91 |
| Penicillium | 4.56 | 0.81 | Chaetomium | 2.77 | 0.64 |
| Pithoascus | 3.94 | 1.78 | Pezizaceae* | 2.61 | 0.6 |
| Botryotrichum | 3.52 | 1.05 | Preussia | 2.03 | 0.61 |
| Aspergillus | 2.79 | 1.61 | Comoclathris | 1.78 | 0.61 |
| Preussia | 2.6 | 1.33 | Fusarium | 1.75 | 0.33 |
| Fusarium | 2.22 | 0.42 | Clarireedia | 1.69 | 0.75 |
| Coniochaeta | 2.12 | 0.82 | Neosetophoma | 1.65 | 0.5 |
| Chaetomium | 1.99 | 0.4 | Neoschizothecium | 1.58 | 0.48 |
| Fusariella | 1.81 | 1.72 | Iodophanus | 1.18 | 0.34 |
| Ascobolus | 1.58 | 0.47 | Aspergillus | 1.16 | 0.42 |
| Acremonium | 1.47 | 0.4 | Phaeosphaeriaceae* | 1.14 | 0.24 |
| Schizothecium | 1.44 | 0.32 | Ascobolus | 1.1 | 0.29 |
| Neocamarosporium | 1.3 | 1.03 | Rutstroemia | 1.1 | 0.41 |
| Neoschizothecium | 1.28 | 0.34 | Aureobasidium | 1.08 | 0.19 |
| Gymnoascus | 1.19 | 0.25 | Schizothecium | 1.03 | 0.3 |
| Stagonosporopsis | 1.14 | 1.07 | Botryotrichum | 1.01 | 0.21 |
| Gibellulopsis | 0.89 | 0.84 | Lecythophora | 0.96 | 0.49 |
| Iodophanus | 0.89 | 0.32 | Naganishia | 0.9 | 0.26 |
| Pseudorobillarda | 0.83 | 0.41 | Pseudogymnoascus | 0.9 | 0.34 |
| Alfaria | 0.73 | 0.63 | Podospora | 0.9 | 0.16 |
| Thelebolus | 0.72 | 0.29 | Apiosordaria | 0.83 | 0.3 |
| Podospora | 0.7 | 0.2 | Mortierella | 0.75 | 0.14 |
| Cladorrhinum | 0.62 | 0.28 | Gymnoascus | 0.69 | 0.16 |
| Pyrenophora | 0.57 | 0.26 | Septoriella | 0.57 | 0.28 |
| Monosporascus | 0.56 | 0.26 | Neostagonospora | 0.55 | 0.26 |
| Septoriella | 0.55 | 0.19 | Pezizomycotina* | 0.55 | 0.15 |

* = Incertae sedis, sem = standard error of the mean
