## Supplementary material for "Urban soils along the Kern River and Los Gatos Creek are hotspots for *Coccidioides* in the San Joaquin Valley of California": S12 Table

**S12 Table:** Indicator species for *Coccidioides* positive and *Coccidioides* negative soil samples collected from within Kern River rodent burrows. Significance was calculated in indicspecies version 1.7.12 with 1000 permutations. Sequences were cross-referenced via NCBI Nucleotide Blast (https://blast.ncbi.nlm.nih.gov/) for each prospective indicator species. • = equally likely alternative species (in subsection). IndVal and p-value apply to all alternative species.

*Coccidioides* positive samples (n = 12)

| | BLASTN % | Reads | IndVal | p-value | |
|---|---|---|---|---|---|
| *Aspergillus salinicola* | 99.2 | 3044 | 0.67 | 0.018 | * |
| *Gymnoascus dugwayensis* | 98.5 | 8128 | 0.849 | 0.012 | * |
| *Coniochaeta discospora* | 100.0 | 6449 | 0.909 | 0.008 | ** |
| *Massariosphaeria roumeguerei* | 95.1 | 516 | 0.69 | 0.025 | * |
| *Montagnea arenaria* | 99.5 | 780 | 0.816 | 0.001 | *** |
| *Myxotrichum deflexum* | 100.0 | 3065 | 0.847 | 0.023 | * |
| *Penicillium citreonigrum* | 100.0 | 847 | 0.56 | 0.022 | * |
| • *Penicillium restrictum* | | | | | |
| • *Penicillium fundyense* | | | | | |
| • *Penicillium toxicarium* | | | | | |
| *Pleuroascus nicholsonii* | 100.0 | 2244 | 0.867 | 0.01 | * |

*Coccidioides* negative samples (n = 16)

| | BLASTN % | Reads | IndVal | p-value | |
|---|---|---|---|---|---|
| *Cladorrhinum flexuosum* | 99.7 | 2969 | 0.773 | 0.037 | * |
| *Fusariella hughesii* | 99.1 | 1182 | 0.783 | 0.022 | * |
| *Fusarium solani* | 100.0 | 969 | 0.658 | 0.031 | * |
| • *Fusarium falciforme* | | | | | |
| • *Fusarium obliquiseptatum* | | | | | |
| • *Umbelopsis nana* | | | | | |
| • *Fusarium oxysporum* | | | | | |

* = $p < 0.05$, ** = $p < 0.01$, *** = $p \leq 0.001$, IndVal = indicator value
