## Supplementary material for "Urban soils along the Kern River and Los Gatos Creek are hotspots for *Coccidioides* in the San Joaquin Valley of California": S13 Table

**S13 Table:** Indicator species for *Coccidioides* positive soil samples collected from within Los Gato Creek rodent burrows. Significance was calculated in indicspecies version 1.7.12 with 1000 permutations. Sequences were cross-referenced via NCBI Nucleotide Blast (https://blast.ncbi.nlm.nih.gov/) for each prospective indicator species. • = equally likely alternative species (in subsection). IndVal and p-value apply to all alternative species.

| | *Coccidioides* positive samples (n = 11) | | | | |
|---|---|---|---|---|---|
| | BLASTN % | Reads | IndVal | p-value | |
| *Alternaria lolii* | 100.0 | 4095 | 0.774 | 0.012 | * |
| *Coniochaeta polymorpha* | 98.4 | 15515 | 0.897 | 0.006 | ** |
| *Kalmusia utahensis* | 100.0 | 151 | 0.739 | 0.019 | * |
| *Monosporascus ibericus* | 96.8 | 577 | 0.739 | 0.017 | * |
| *Preussia polymorpha* | 98.1 | 6910 | 0.888 | 0.048 | * |
| *Sporormiella megalospora* | 98.1 | | | | |
| *Preussia terricola* | 98.1 | | | | |

* = $p < 0.05$, ** = $p < 0.01$, *** = $p \leq 0.001$, IndVal = indicator value
