## Supplementary material for "Urban soils along the Kern River and Los Gatos Creek are hotspots for *Coccidioides* in the San Joaquin Valley of California": S14 Table

**S14 Table:** Indicator species for *Coccidioides* negative soil samples collected from within Los Gatos Creek rodent burrows. Significance was calculated in indicspecies version 1.7.12 with 1000 permutations. Sequences were cross-referenced via NCBI Nucleotide Blast (https://blast.ncbi.nlm.nih.gov/) for each prospective indicator species. • = equally likely alternative species (in subsection). IndVal and p-value apply to all alternative species.

| *Coccidioides* negative samples (n = 9) | | | | | | |
|---|---|---|---|---|---|---|
| | | BLASTN % | Reads | IndVal | p-value | |
| *Aspergillus cristatus*<br>• *Aspergillus amstelodami*<br>• *Aspergillus chevalieri*<br>• *Aspergillus medius*<br>• *Aspergillus ruber*<br>• *Aspergillus costiformis* | • *Aspergillus intermedius*<br>• *Aspergillus montevidensis*<br>• *Aspergillus medius*<br>• *Aspergillus heterocaryoticus*<br>• *Aspergillus caperatus* | 100.0 | 331 | 0.745 | 0.009 | ** |
| *Chaetomium madrasense*<br>• *Chaetomium contagiosum*<br>• *Chaetomium grande*<br>• *Chaetomium interruptum* | • *Chaetomium megalocarpum*<br>• *Chaetomium ascotrichoides*<br>• *Chaetomium globosum* | 100.0 | 3542 | 0.79 | 0.007 | ** |
| *Circinella muscae* | • *Fennellomyces linderi* | 100.0 | 949 | 0.739 | 0.033 | * |
| *Holtermanniella festucosa*<br>• *Holtermanniella wattica* | • *Holtermanniella takashimae*<br>• *Holtermanniella festucosa* | 99.4 | 482 | 0.899 | 0.003 | ** |
| *Dactylonectria torresensis* | • *Dactylonectria macrodidyma* | 100.0 | 180 | 0.732 | 0.028 | * |
| *Enterocarpus grenotii* | • *Basipetospora chlamydospora* | 99.1 | 1795 | 0.822 | 0.014 | * |
| *Ustilago bullata* | | 100.0 | 169 | 0.786 | 0.009 | ** |
| *Fusarium oxysporum*<br>• *Fusarium solani*<br>• *Fusarium foetens* | • *Fusarium graminearum*<br>• *Fusarium fujikuroi* | 100.0 | 549 | 0.667 | 0.026 | * |
| *Mortierella alpina* | | 100.0 | 577 | 0.808 | 0.013 | * |
| *Linnemannia gamsii* | | 98.5 | 441 | 0.717 | 0.038 | * |
| *Myrmecridium schulzeri* | | 100.0 | 680 | 0.751 | 0.046 | * |
| *Paraphoma fimeti* | • *Septoriella neoarundinis* | 100.0 | 365 | 0.803 | 0.004 | ** |
| *Pseudoechria decidua* | | 96.2 | 173 | 0.745 | 0.008 | ** |
| *Solicoccozyma aeria* | | 99.7 | 1189 | 0.814 | 0.011 | * |
| *Phoma sojicola*<br>• *Ascochyta rabiei*<br>• *Nothophoma quercina*<br>• *Stagonosporopsis* | • *Microsphaeropsis olivacea*<br>• *Allophoma zantedeschiae*<br>• *Allophoma labilis*<br>• *Ascochyta medicaginicola* | 99.3 | 219 | 0.667 | 0.03 | * |
| *Tetracladium furcatum* | • *Tetracladium setigerum* | 99.6 | 157 | 0.725 | 0.029 | * |

* = $p < 0.05$, ** = $p < 0.01$, *** = $p \leq 0.001$, IndVal = indicator value
