## Supplementary material for "Urban soils along the Kern River and Los Gatos Creek are hotspots for *Coccidioides* in the San Joaquin Valley of California": S15 Table

**S15 Table:** Indicator species for *Coccidioides* samples (positive and negative) collected from surface soils outside Los Gatos Creek rodent burrows. Significance was calculated in indicspecies version 1.7.12 with 1000 permutations. Sequences were cross-referenced via NCBI Nucleotide Blast (https://blast.ncbi.nlm.nih.gov/) for each prospective indicator species. • = equally likely alternative species (in subsection). IndVal and p-value apply to all alternative species.

*Coccidioides* positive samples (n = 10)

| | BLASTN % | Reads | IndVal | p-value | |
|---|---|---|---|---|---|
| *Coniosporium apollinis* | 100.0 | 4420 | 0.91 | 0.014 | * |
| *Coprinopsis gonophylla* | 99.7 | 2394 | 0.831 | 0.02 | * |
| *Mycotypha microspora* | 100.0 | 155 | 0.747 | 0.029 | * |

*Coccidioides* negative samples (n = 10)

| | BLASTN % | Reads | IndVal | p-value | |
|---|---|---|---|---|---|
| *Dothiora viticola* | 99.4 | 487 | 0.775 | 0.008 | ** |
| *Podospora vesticola* | 99.0 | 3908 | 0.768 | 0.023 | * |
| • *Schizothecium carpinicola* | | | | | |
| *Ramimonilia apicalis* | 97.6 | 536 | 0.759 | 0.042 | * |

* = p < 0.05, ** = p < 0.01, *** = p ≤ 0.001, IndVal = indicator value
